# Characterizing 3D RNA structural features from DMS reactivity

**DOI:** 10.1101/2024.11.21.624766

**Authors:** D. H. Sanduni Deenalattha, Chris P. Jurich, Bret Lange, Darren Armstrong, Kaitlyn Nein, Joseph D. Yesselman

**Affiliations:** Department of Chemistry, University of Nebraska, 639 North 12th St, Lincoln, NE 68588, USA

## Abstract

Dimethyl sulfate (DMS) chemical mapping probes RNA structure, where low reactivity is generally interpreted as Watson-Crick (WC) base pairs and high reactivity as unpaired nucleotides. Studies examining DMS reactivity of RNAs with known 3D structures have identified nucleotides that deviate from this interpretation with distinct solvent accessibility and hydrogen bonding patterns. Understanding the frequency of these outliers and their recurring structural 3D features remains incomplete. To address this knowledge gap, we systematically analyzed DMS reactivity patterns across a library of 7,500 RNA constructs containing two-way junctions with known 3D structures. We observe DMS reactivity exists on a continuum over four orders of magnitude with approximately 10% overlap in reactivity between WC and non-WC nucleotides. We find that non-WC bases with WC-like DMS protection exhibit increased hydrogen bonding and decreased solvent accessibility, whereas WC pairs exhibiting greater DMS reactivity tend to flank junctions, correlating with weaker base stacking and greater junction dynamics. Furthermore, we discover that DMS reactivity values in non-canonical pairs correlate with atomic distances and base pair geometry, enabling discrimination between different 3D conformations. These DMS reactivity patterns indicate that DMS reactivity provides atomic-scale information about RNA 3D conformations, which can be used to model RNA structures and dynamics.

## Introduction

Structured RNAs are pivotal in fundamental biological processes, including protein translation, mRNA maturation, and telomere maintenance (1-3). To perform these functions, many RNAs fold into intricate secondary and tertiary structures that often undergo conformational changes in response to stimuli (4-8). Elucidating these functions requires knowledge of RNA folding and conformational dynamics.

While high-resolution 3D structures provide valuable atomic-level insights, they capture static snapshots, which may fail to elucidate conformational transitions and their associated energetics crucial for RNA function (9, 10). Chemical mapping offers an orthogonal and complementary approach to high-resolution structure determination. These approaches employ small molecule reagents that chemically modify nucleotides based on their local environment, giving insights into dynamic conformation changes and thermodynamics (11-24). In particular, dimethyl sulfate (DMS) can provide information about RNAs in and out of their cellular contexts and its use has accelerated due the ability to read out modifications via next-generation sequencing (24, 25). These next-generation sequencing approaches can read out DMS methylations at the N1 position of adenine and the N3 position of cytosine, typically leaving nucleotides involved in Watson-Crick (WC) base pairs unmodified. These specific modification patterns have enabled both validation of RNA structures predicted through phylogenetic or thermodynamic approaches and generation of new structural models (24, 26-30). Furthermore, analysis of RNA sequences that contain multiple DMS modifications in a single read has enabled the computational separation of the data into clusters that reveal distinct RNA secondary structures (31-35).

Recently we demonstrated that there is a direct relationship between DMS reactivity values and thermodynamics of tertiary contact formation (13). This ability to provide quantitative information about RNA 3D structure inspired us to confront two critical limitations in current DMS approaches. First, current RNA structure prediction methods use DMS reactivity values to bias folding algorithms through pseudo-energy terms, where lower reactivity increases the likelihood of Watson-Crick base pairing and higher reactivity decreases it (24, 26-30). While these techniques have been successful in building secondary structure models this simplified interpretation discards potentially valuable quantitative information about modification frequencies that report on 3D structural features. Second, over the past decades there is a wealth of studies that indicate there are exceptions to the general interpretation that low reactivity implies WC pairs and high reactivity are non-WC residues when compared to high resolutions structures. Furthermore, there are cases such as sheared G-A pairs where the A is hyper reactive, displaying reactivity higher than residues that are fully exposed to solvent (12, 20, 36-39). The structural mechanisms driving these effects stem from variations in solvent accessibility and hydrogen bonding patterns (40-43). However, the frequency of occurrence, the precise 3D structural features that generate them, and their identification from DMS reactivity data remain unknown.

In this study, we designed a comprehensive RNA library to establish quantitative relationships between DMS reactivity and RNA 3D structure. The library comprised 7,500 RNA constructs containing distinct two-way junctions with known 3D structures. DMS chemical mapping of these constructs revealed reactivity patterns across diverse sequence contexts. Analysis of the mapping data demonstrated DMS reactivity exists as a continuous distribution with substantial overlap between WC and non-WC nucleotides. The data revealed correlations between DMS reactivity and junction asymmetry, base-stacking interactions, and neighboring base pair sequences. Non-canonical base pairs exhibited characteristic reactivity patterns that correspond to specific 3D conformations, enabling structural feature identification from DMS data. The systematic relationships between DMS reactivity and RNA 3D structural features will provide a foundation for interpreting chemical mapping data of functional RNAs and determining their conformational states.

## Results

### Designing a massive library to quantitatively relate DMS reactivity to RNA structure

To build a quantitative relationship between RNA structure and DMS reactivity, we developed a systematic approach using RNA elements with known 3D structures. We extracted two-way junctions from the RNA non-redundant database (44), which are non-WC interactions flanked by two WC base pairs. These junctions are ideal for our study because they maintain their structure when isolated from larger RNAs (4, 45-47). They are fundamental building blocks in functional RNAs, playing critical roles in ligand binding and catalysis (48-50). We found 177 unique RNA two-way junctions that were isolatable, i.e., had no more than two hydrogen bonds to non-motif residues (**See methods**). These junctions represent diverse RNA structural elements, including kink turns (51), sarcin-ricin loops (52), bulges, and internal loops (**Supplemental Figure S1, Supplemental Table S1**). We supplemented our dataset with 536 1×1 and 2×2 symmetrical junctions without known 3D structures (**Supplemental Table S2**). Previous work has shown that these small, symmetric junctions often comprise non-canonical base pairs (53). These additional junctions expand our ability to systematically observe trends among different types of potential non-canonical pairs.

We engineered a massive RNA library by incorporating these junctions into 7,500 unique RNA constructs. Each construct was designed as a 150-nucleotide sequence containing 5-7 junctions arranged within stable hairpin structures (**Figure 1A**). This hairpin architecture was crucial - providing a stable structural scaffold ensured each junction would fold into its intended conformation rather than forming alternative structures (**see Methods**). Each junction appears 30 times on average, ranging from 5 to 104 occurrences (**Figure 1E**). This redundancy enables the calculation of average DMS reactivity per junction and reveals how local sequence context influences junction reactivity.

**Figure 1:**
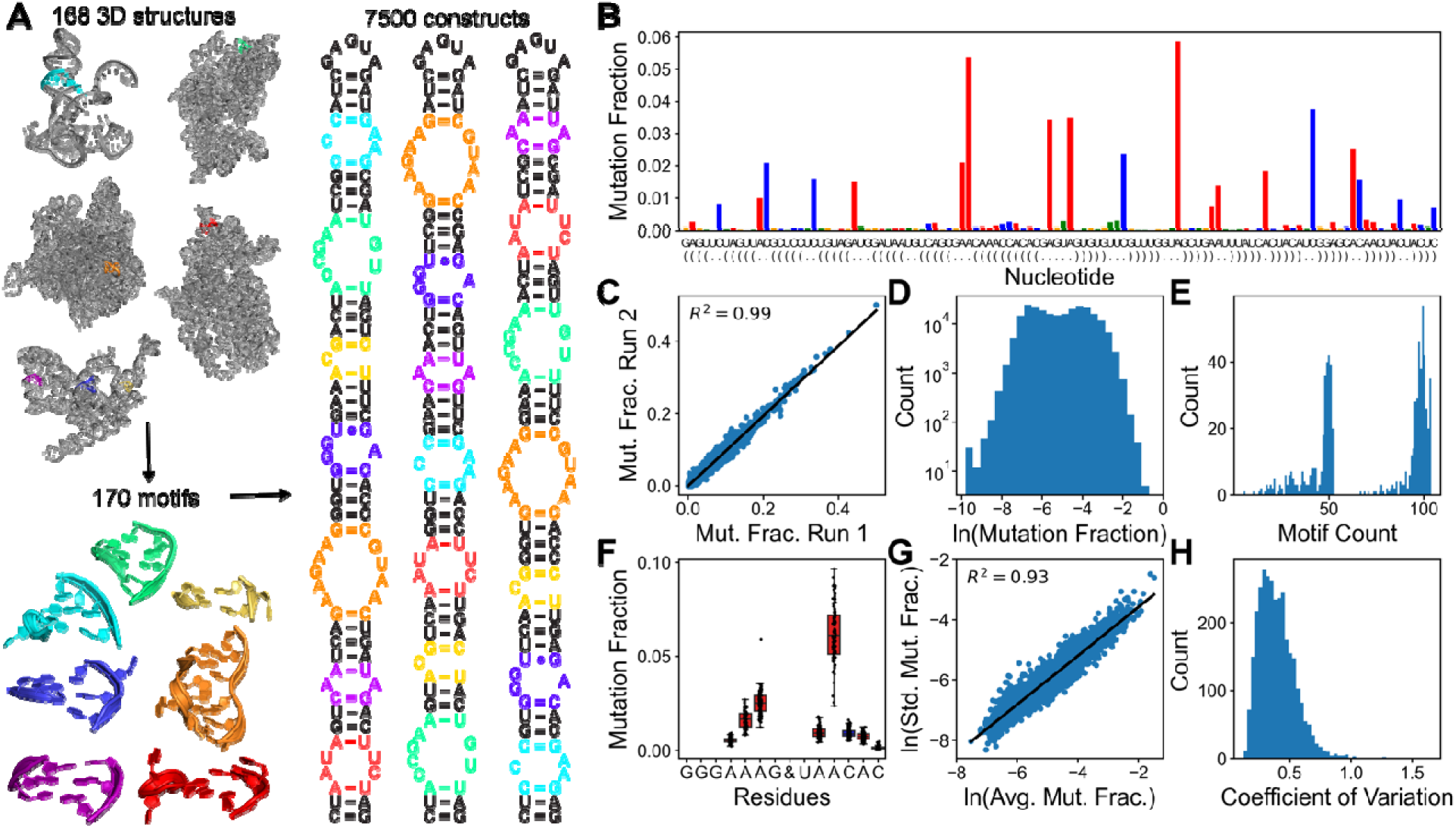
Design and validation of a large-scale RNA library for DMS structure-reactivit analysis. (A) Library design strategy: Isolatable RNA motifs were extracted from 3D structures an combined with engineered variants to create stable hairpin constructs containing multiple motif separated by helices. (B) A representative construct shows the secondary structure and DM reactivity data. Nucleotides are color-coded: adenine (red), cytosine (blue), uracil (green), an guanine (gold). The height of the bars indicates DMS reactivity. (C) Reproducibility of DM measurements across independent experiments. Each point represents a nucleotide’ mutational fraction (R² = 0.99, n = 240,000 measurements). (D) Distribution of DMS reactivit values shown on a natural logarithmic scale, spanning four orders of magnitude (6.0 x 10⁻L t 0.5). (E) Frequency distribution of motif occurrences within the library. (F) Example of reactivit consistency: The motif "GGGAAAG&UAACAC" with secondary structure “(…..(&)…..)” show similar DMS reactivity patterns across multiple sequence contexts, demonstrating measurement reproducibility. (G) Measurement variability analysis: Coefficient of variation (CV = standard deviation/mean) for each nucleotide position across all motif instances. (H) Impact of structural context: Comparison of CV distributions when nucleotides are grouped by second flanking pair identity versus ungrouped, showing reduced variability with grouping (see **Supplemental Figure 4** to compare to the random grouping of the same size).

### DMS reactivity is highly reproducible and is primarily governed by local sequence and structure

We performed DMS mutational profiling with sequencing (DMS-MaPseq) on our library of 7500 constructs, with a high average read depth of 38,000 reads per sequence (**Figure 1B, Supplemental Figure S2**). Our measurements revealed DMS reactivities spanning four orders of magnitude (6.0 x 10^-5^ to 0.5). Based on previous research, we employed the natural logarithm of DMS reactivity, which allows for a more intuitive interpretation of the data while preserving the full range of observed reactivities (**Figure 1D**) (13). Replicate experiments showed excellent reproducibility (R² = 0.99) for the 240,000 DMS measurements (**Figure 1C**), though the correlation decreased with reactivity values below 0.001 (R² = 0.37; **Supplemental Figure S3**). This lower bound corresponds to our no-modification background mutation rate of 0.0014.

To quantify how consistently each junction behaved across different sequence contexts, we calculated the coefficient of variation (CV) - a standardized measure of variability that divides standard deviation by the mean. A low CV would indicate that a junction’s DMS reactivity remains constant regardless of its position in different constructs. In contrast, a high CV would suggest that surrounding sequences strongly influence its reactivity. We found that the average CV was 0.36 across all nucleotides (**Figure 1H**), with WC pairs showing more variability (CV = 0.42) than non-WC residues (CV = 0.30). The elevated CV value for WC pairs is due to the lower DMS reactivity values which will generally have more affected by the number of reads. CV values between 0.2 and 0.3 are usually considered moderate and above 0.3 is high. Which lead us to consider other structural effects to explain the CV values.

To further investigate the cause of the size of the average DMS reactivity CV, we analyzed the effects of the next WC pair after the flanking pair (the second flanking pair). When we grouped our data based on the second flanking pair identity, the average CV for WC pairs decreased from 0.42 to 0.34, and non-WC residues reduced from 0.30 to 0.22. To ensure these reductions weren’t simply an artifact of dividing our data into smaller groups, we performed a control analysis using random groupings of the same size, which showed significantly smaller CV reductions, 0.37 for WC pairs and 0.27 for non-WC residues (**Supplemental Figure S4**). These data indicate that DMS reactivity is reproducible over an extensive range of values and largely depends on local effects including the sequence of junction and its neighboring base pairs.

### DMS reactivity values are continuous, and a significant overlap exists between Watson-Crick and non-Watson-Crick nucleotides

A key purpose of this study is to systematically investigate the relationship between high-resolution structure features and DMS reactivity. Most RNA structure prediction methods use a pseudo-free energy term that reduces likelihood of paired based on how high the reactivity of a nucleotide is. Based on this assumption, there should be two distinct populations of DMS reactivity values in our dataset, one with low reactivity that is WC pairs and the other higher for non-WC residues. However, comparing the DMS reactivity of flanking WC pairs with non-WC residues revealed overlapping distributions (**Figure 2A**) (including non-flanking WC pairs gives similar distributions and analysis but lack 3D structural information see **Supplemental Figure S5**).

**Figure 2:**
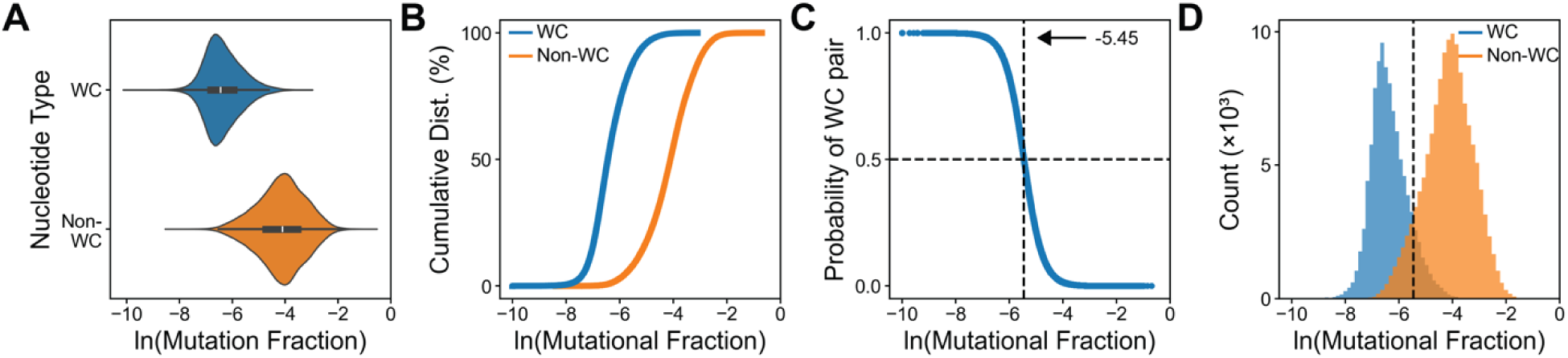
Quantitative analysis of DMS reactivity in flanking Watson-Crick and non-Watson-Crick nucleotides. (A) Reactivity distributions reveal a significant overlap between nucleotides in flanking WC pairs (blue) and non-WC positions (orange), challenging the simple binary interpretation of DMS data. (B) Cumulative reactivity distributions comparing WC paired versus non-WC nucleotides as a function of the natural log of the mutational histogram. (C) Logistic regression analysis establishing the probability of WC pairing based on DMS reactivity. The horizontal dashed line marks the 50% probability threshold, corresponding to a natural log mutation fraction of -5.45 (mutation fraction = 0.0043). (D) Distribution of nucleotides relative to the 50% probability threshold. While this cutoff optimally separates WC from non-WC nucleotides, significant overlap demonstrates the limitations of using fixed reactivity thresholds for structure prediction.

We applied a logistical regression to find the best reactivity cutoff to quantitatively distinguish WC pairs from non-WC residues based on DMS reactivity. The analysis identified a DMS reactivity threshold of 0.0043 (ln (0.0043) = -5.45), corresponding to a 50% probability of a nucleotide being in a WC pair (**Figure 2C**). Using this threshold value, we found that 9.87% of non-WC residues (11,597/117,498) showed low reactivity typical of WC pairs, while 8.92% of WC residues (10,399/116,579) displayed unexpectedly high reactivity.

### DMS reactivity of flanking WC pairs report on sequence, structure, and dynamics

To characterize the structural features driving DMS reactivity in flanking WC base pairs, we analyzed conformational, and sequence elements associated with high reactivity (reactivity > 0.0043) in these canonical pairs. We first observed a significant difference in the frequency of high reactivity flanking pairs between C-G (1%) and A-U (19%) pairs (**Figure 3A**). This substantial difference cannot be attributed to sampling bias, as our dataset contained comparable numbers of A-U and C-G pairs (47,652 and 52,121, respectively). This significant difference reflects the C-G pair’s greater thermodynamic stability, providing greater protection against DMS modification than A-U pairs. These data indicate that base pair identity plays a role in understanding the difference in DMS reactivity of flanking WC pairs.

**Figure 3:**
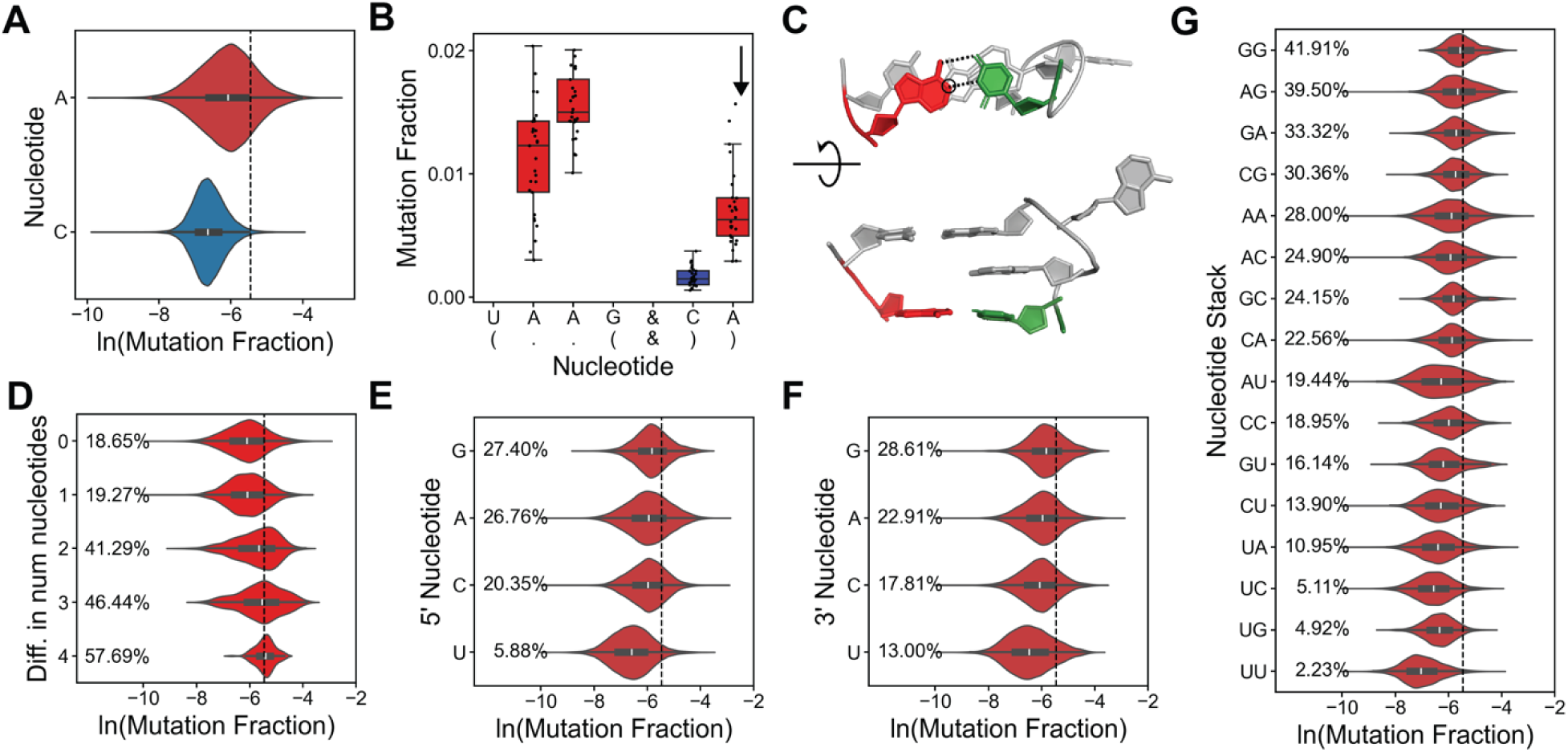
Sequence context and structural features influence Watson-Crick pair reactivity. (A) DMS reactivity distributions comparing adenines in A-U pairs versus cytosines in C-G pairs. The vertical dotted line (natural log of reactivity = -5.45) is the 50% likelihood of being a WC base pair. A-U pairs are frequently more reactive than C-G pairs. (B-C) Example of WC pair that are highly reactive but maintain ideal geometry. (D) Distributions of the natural log of reactivity for As in A-U flanking pairs as a function of the asymmetry of the non-WC paired residues where 0 is symmetrical, and 4 is four more residues on one side than the other. (E) The distribution of the natural log of reactivity as a function of the 5′ residue or the residue that appears right before the A in the A-U pair. (F) The distribution of the natural log of reactivity as a function of the 3′ residue or the residue that appears right after the A in the A-U pair. (G) The combined influence of flanking sequence context. Two-nucleotide patterns (e.g., "GG" = 5′-GAG-3′) reveal strong neighboring effects, with purine-rich contexts promoting higher reactivity. Similar trends were observed for cytosines (**Supplemental Figure S17**).

To examine whether local structural distortions in WC pairs could account for reactivity differences, we examined each motif’s high-resolution structure. Analysis of 306 high-resolution structures revealed no correlation between DMS reactivity and base pair geometric parameters (shear, stretch, stagger, buckle, propeller, and opening) or overall deviation from ideal geometry (**Figure 3B-C, Supplemental Figure S6**). Further analysis by specific base pair types (A-U, U-A, G-C, or C-G) revealed no correlation improvement. These findings indicate that the static, lowest energy conformations of flanking WC pairs, as captured in high-resolution crystal structures, do not provide sufficient information to explain the observed variations in DMS reactivity.

Given the lack of correlation between DMS reactivity and static structural features, we explored the role of RNA dynamics in flanking base pair reactivity. Previous studies suggest symmetric junctions form stable non-canonical pairs, while asymmetric junctions exhibit increased flexibility (53). We quantified this relationship by analyzing junction asymmetry, which is defined by the difference in residue numbers on each side (0 for symmetric to 4 for highly asymmetric). Our analysis revealed a correlation between junction asymmetry and elevated DMS reactivity in flanking pairs (**Figure 3D**). This pattern aligns with known dynamic structures like the 3×0 HIV-1 TAR bulge, where the flanking AU pair forms transiently (54).

We also analyzed how local sequence context influences flanking pair reactivity. Examining residues adjacent to flanking pairs revealed sequence-dependent patterns of DMS reactivity (**Figure 3E-G**). We found purines at either 5′ or 3′ positions significantly increased the probability of high reactivity in flanking WC pairs. These effects compound flanking pairs with guanines on both sides showed a 40% probability of elevated reactivity compared to only 2% for two uracils. These sequence-dependent reactivity patterns suggest a balance between base stacking and hydrogen bonding in RNA structure. Purine-rich environments favor stacking interactions over hydrogen bonding, increasing flanking pair flexibility and DMS reactivity. This model aligns with multiple high-resolution structures, where stacked purines forgo hydrogen bonding (**Supplemental Figure S7**), suggesting a general principle in RNA structural organization.

### Non-canonical base pairs protect against DMS modification through hydrogen bonding and decreased solvent accessibility

To characterize structural features causing low DMS reactivity in non-WC nucleotides, we analyzed cases where non-canonical interactions exhibited WC-like protection. To enhance RNA structure prediction accuracy from DMS data, we identified cases where non-canonical interactions could be misinterpreted as WC pairs due to low reactivity. Analysis of nucleotides with known 3D structures revealed that 11% of non-canonical pairs showed reactivity below our WC threshold (< 0.0043), compared to only 2.5% of unpaired nucleotides. This 4.4-fold difference indicates non-canonical pairing can protect nucleotides from DMS modification similarly to WC pairs (**Figure 4A-B**). Our engineered 1x1 and 2x2 motifs without known structures confirmed this pattern, where 14.50% and 9% of potential non-canonical pairs, respectively, showed low reactivity. These results indicate that non-canonical interactions frequently generate WC-like protection patterns.

**Figure 4:**
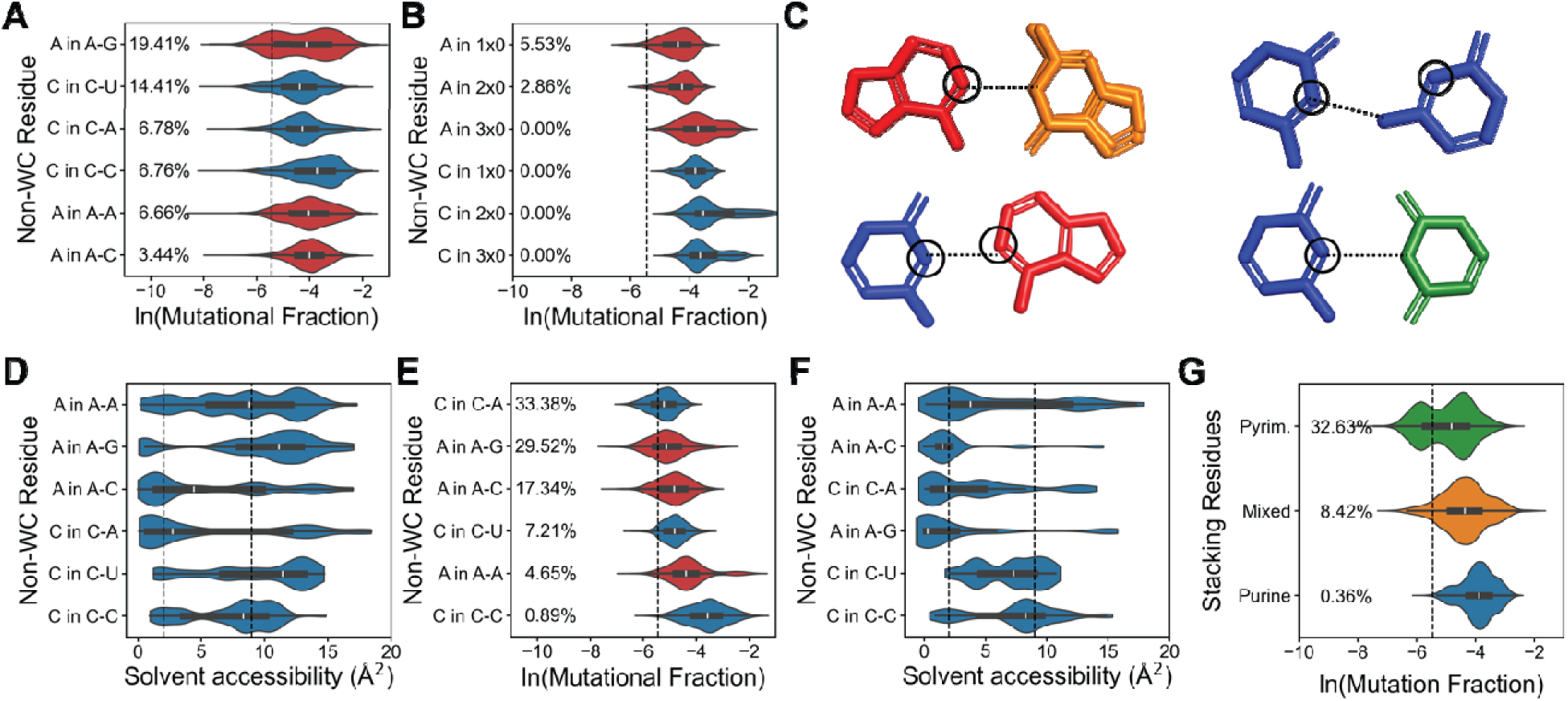
Structural and sequence determinants of low reactivity in non-canonical pairs. (A) DMS reactivity distributions for non-canonical pairs with known structures. A vertical dotte line indicates the 50% WC probability threshold (-5.45). Nucleotides left of this line exhibit WC-like protection. (B) Reactivity distribution of unpaired nucleotides in bulges, providing baselin comparison for non-canonical pairs. (C) Representative examples of conformations of low-reactivity non-canonical pairs: A-G, A-C, C-C, and C-U, colored as A (red), C (blue), G (orange), and U (green). Each illustrates how specific hydrogen bonding patterns can protect DM modification sites. (D) Solvent accessibility of DMS modification sites (adenine N1, cytosine N3) across non-canonical pairs. Horizontal lines indicate average accessibility for WC pairs (2 A) and unpaired nucleotides (upper), demonstrating a correlation between accessibility an reactivity. (E) DMS reactivity distributions for non-canonical pairs with known structures in 1x mismatches. A vertical dotted line indicates the 50% WC probability threshold (-5.45). Nucleotides left of this line exhibit WC-like protection. (F) Solvent accessibility analysis focuse on 1×1 mismatches, revealing distinct patterns of nucleotide protection in symmetric contexts. (G) The impact of neighboring sequences on C-U mismatch reactivity shows how local context modulates DMS accessibility in non-canonical pairs.

We analyzed reactivity patterns across mismatched pairs (A-A, A-G, A-C, C-C, C-U) using structures with known 3D conformations. Low reactivity frequencies varied by pair type: A-G (19.40 %), C in C-U (14.39 %), C in C-A (6.78 %), C-C (6.75 %), A-A (6.65 %), and A in A-C (3.44 %) (**Figure 4A**). This protection stems from hydrogen bonding at DMS modification sites (N1 of adenine, N3 of cytosine). For example, A-G pairs often form cis Watson-Crick/Watson-Crick (cWW) arrangements where stable a N1-N1 hydrogen bond provide protection. Similarly, low-reactivity C-A pairs typically show cWW conformations where cytosine’s N3 hydrogen bonds to adenine’s N1. These conformations must involve a protonation of one nucleotide, consistent with previously by NMR (55-58) (**Figure 4C**).

Solvent accessibility plays a crucial role in DMS reactivity. In WC pairs, the N1/N3 positions are shielded from solvent, reducing DMS modification. Analysis of 695 nucleotides from high-resolution structures revealed a moderate correlation (R² = 0.41) between DMS reactivity and solvent accessibility of these modification sites (**Supplemental Figure S8**). This relationship extends to non-canonical pairs – those with lower DMS reactivity showed reduced solvent accessibility at N1 and N3 atoms (**Figure 4D**). This pattern is most evident in 1x1 mismatches, where 33.38% of cytosines in C-A pairs and 29.52% of adenines in A-G pairs demonstrated WC-like reactivity (**Figure 4E**). These nucleotides had low solvent accessibility, with 51.66% of C-A pairs and 69.25% of A-G pairs showing values below 2 Å^2^, typical of WC pairs (**Figure 4F**, **Supplemental Table S3**). These findings provide strong evidence that the DMS reactivity of non-canonical pairs can approach that of WC pairs consistent with the reduced solvent accessibility of the target atoms.

The local stacking environment significantly influences non-canonical pair reactivity. While adenines in A-A, A-G, and A-C pairs showed minimal stacking effects, cytosines in C-A, C-C, and C-U pairs were strongly influenced by their neighboring bases (**Supplemental Figure S9**). Pyrimidines flanking these cytosine-containing pairs correlated with reduced reactivity. This effect was particularly dramatic in C-U pairs – 32.31% of cytosines with pyrimidine neighbors showed WC-like reactivity, compared to just 0.35% when flanked by purines (**Figure 4G**). This hundred-fold difference likely results from competing structural forces. In pyrimidine-rich environments, cytosine’s weak stacking ability favors hydrogen bond formation with its partner, shielding the N3 position from DMS. Conversely, purine stacking may disrupt hydrogen bonding, exposing the N3 position and increasing DMS reactivity.

### Non-canonical base pairs have distinct reactivity relationships that report 3D structure features

We explored the potential of DMS reactivity data to reveal 3D structural information about non-canonical base pairs. We found that A-G, C-A, and C-C reactivity patterns correlate with specific atomic distances, providing insights into base-pair conformations (**See Supplemental Figure S10** for other pairs with weaker correlations). For A-G pairs, we found a correlation (R² = 0.51, n = 122) between the phosphate-to-phosphate (P-P) distance and adenine reactivity (**Figure 5A**). This correlation resulted from the longest P-P distance in cis Watson-Crick/Watson-Crick (cWW) conformations, corresponding to the lowest reactivity values. In contrast, shorter P-P distances were associated with trans Sugar/Hoogsteen (tSH) conformations and the highest reactivity values (**Figure 5B**). This pattern suggests that DMS reactivity is sensitive to the overall geometry of the base pair. Further analysis revealed that grouping reactivities by base pair conformation yielded distinct clusters, indicating that the specific interaction type is a primary determinant of reactivity patterns (**Figure 5C**). While the P-P distance provided valuable insights, it represents just one of several atomic measurements correlating with base-pairing modes (**Supplemental Figure S11**).

**Figure 5:**
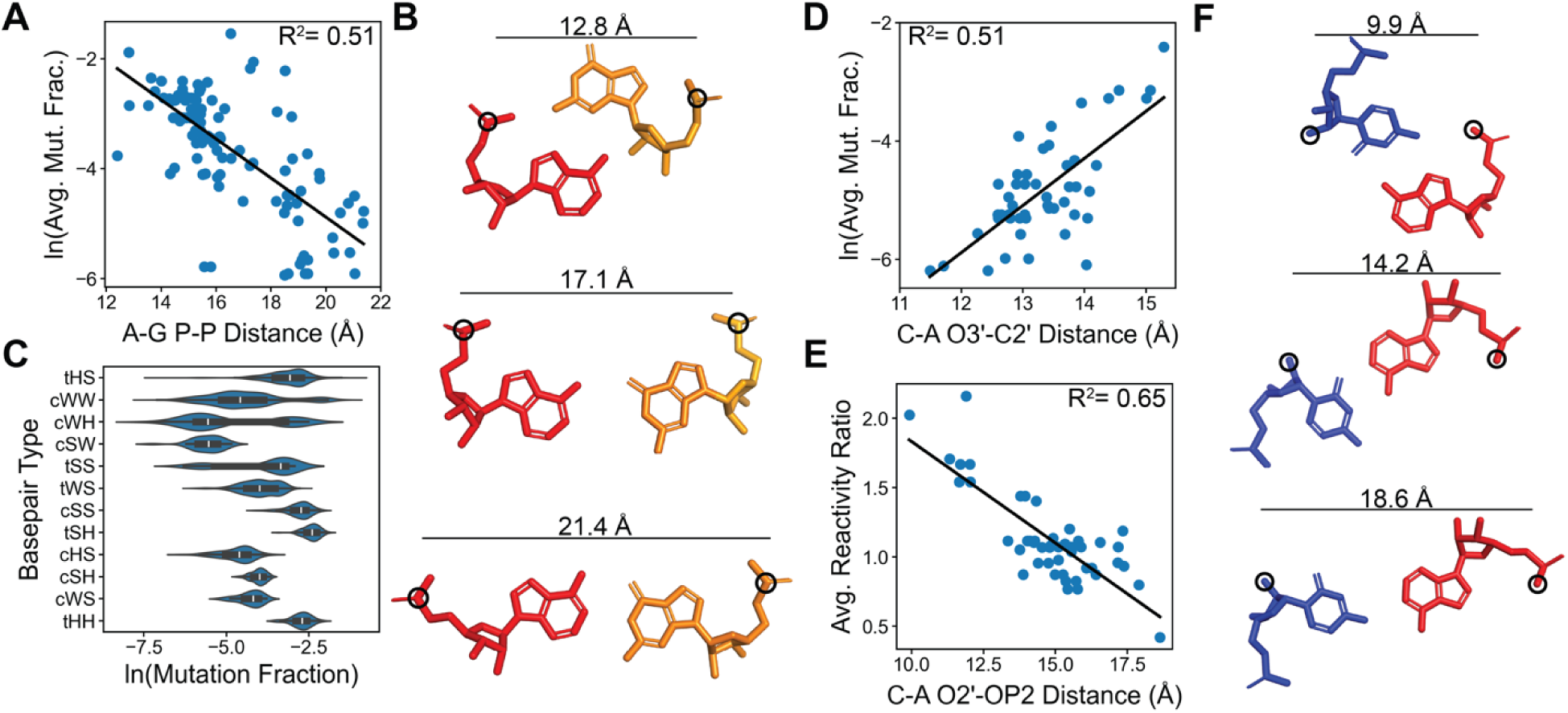
DMS reactivity correlates with RNA 3D structural features of non-canonical pairs. DMS features DMS reactivity analysis reveals quantitative relationships with three-dimensional structural parameters in non-canonical base pairs. (A) The natural log of adenine reactivity in A-G pairs correlates with phosphate-phosphate distance (R² = 0.51, n=122). (B) Representative A-G pairs show short, medium, and long P-P distances. (C) Distribution of adenine reactivity in A-G pairs by base pair conformation type, showing distinct patterns. (D) Cytosine reactivity in C-A pairs correlates with O3′-C2′ distance (R² = 0.51, n=48). (E) The cytosine-to-adenine reactivity ratio in C-A pairs versus O2′-OP2 distance reveals geometric relationships (R² = 0.65, n=48). (F) Representative C-A pairs show short, medium, and long O2′-OP2 distances.

C-A pairs showed multiple correlations between atomic distances and reactivity. The strongest correlation (R² = 0.51, n = 48) was between cytosine’s O3′ atom and adenine’s C2′ atom (**Figure 5D, Supplemental Figure S12**). Short O3′-C2′ distances corresponded to low cytosine reactivity in trans Sugar/Hoogsteen (tSH) and some cis Watson-Crick/Watson-Crick (cWW) conformations. Longer distances showed higher reactivity in cis Watson-Crick/Hoogsteen (cWH) and trans-Watson-Crick/Hoogsteen (tWH) arrangements. An even stronger correlation (R² = 0.65, n = 48) appeared when comparing the ratio of cytosine-to-adenine reactivity with the distance between cytosine’s O2′ and adenine’s OP2 atoms (**Figure 5E-F, Supplemental Figure 13**). This ratio effectively measures the relative solvent accessibility of the modification sites. Ratios above 1 indicate better protection of cytosine’s N3 compared to adenine’s N1, typical in tWH conformations. Ratios near 1 suggest equal accessibility, common in cWW arrangements, while ratios below 1 show better protection of adenine’s N1.

For C-C pairs showed the strongest correlation (R² = 0.64, n = 35) between O3′ and OP2 distances, though with limited samples. Notably, this correlation reveals structural asymmetry, with indicating hydrogen bonding between one cytosine’s O3′ and the other’s OP2 (**Supplemental Figure S14-15).** Together, these correlations between DMS reactivity and atomic distances indicate that chemical mapping data encodes 3D structural information of non-canonical pairs.

## Discussion

In this study we analyzed the DMS reactivity patterns in RNA 3D structures using 7,500 RNA constructs containing known structural motifs and engineered symmetrical junctions. These constructs combine known 3D structural motifs with engineered 1×1 and 2×2 symmetrical junctions, providing data for hundreds of instances of each structural element across different sequence contexts. Results from this library demonstrate that DMS reactivity extends across four orders of magnitude in a continuous distribution. The DMS reactivity patterns remained consistent when motifs were placed in different sequence contexts, although neighboring WC pairs influenced reactivity of both flanking pairs and non-canonical interactions. Controlling for second flanking pair identity reduced DMS reactivity variability and increased reproducibility. Systematic investigation of second and third flanking pair effects could enhance thermodynamic parameters for RNA structure prediction.

Modern RNA structure prediction methods incorporate DMS reactivity through pseudo-energy terms in folding algorithms. These terms favor WC base pairing at positions with low reactivity and disfavor pairing at positions with high reactivity. Prior research identified cases where non-Watson-Crick residues exhibit low reactivity and WC pairs show high reactivity in specific RNA structures. This study quantified the frequency of DMS reactivity patterns that deviate from the canonical model. The analysis revealed 10% of nucleotides diverge from expected patterns, with Watson-Crick pairs showing higher reactivity and non-Watson-Crick residues displaying lower reactivity than predicted. However, these outliers should not be interpreted as errors but as potential indicators of 3D structural features.

Multiple structural features influence reactivity patterns that generate DMS outliers. Watson-Crick pair reactivity depends on base pair composition, with C-G pairs exhibiting five-fold lower reactivity than A-U pairs. Flanking purines and junction asymmetry correlate with increased reactivity. Non-Watson-Crick residues display reduced reactivity through hydrogen bonding patterns. Overall, 11% of non-canonical residues show Watson-Crick protection levels, with specific interactions like A-G pairs reaching 20% protection frequency. These non-canonical geometries restrict solvent accessibility to levels comparable to Watson-Crick pairs. Furthermore, base stacking environments influence reactivity in a position-dependent manner, exemplified by neighboring pyrimidines providing 100-fold greater protection to C-U pairs compared to neighboring purines. This protection derives from competing structural forces - pyrimidines stack poorly, enabling hydrogen bond formation that shields nucleotides from DMS reactivity.

Finally, DMS reactivity patterns provide information for RNA tertiary structure assignment through correlations between reactivity and three-dimensional features. The phosphate-to-phosphate distance in A-G pairs correlates with adenine reactivity (R² = 0.51), enabling discrimination between base-pairing geometries. Additional non-canonical pairs display characteristic correlations between atomic distances and reactivity ratios that identify distinct base-pairing modes, as demonstrated in C-A pairs.

Overall, our findings indicate that DMS chemical mapping data contains more structural information than previously utilized. Future work can build on these relationships between reactivity patterns and structural features to develop more accurate RNA structure prediction methods, particularly for complex structural elements containing non-canonical pairs. These correlations can enable modeling of complex structural elements containing non-canonical pairs and guide tertiary structure determination of RNA motifs with non-canonical interactions.

## Data, materials, and software availability

All data, materials, and software used in this study are available. Unprocessed FASTQ files have been deposited to the Sequence Read Archive (SRA) under the accession PRJNA1188187. All other data is available on Fig Share (10.6084/m9.figshare.27880434). All code used in this study is available on GitHub: https://github.com/YesselmanLabPublications/2025_char_3d_struct_features

## Acknowledgments

This work was supported by the NIH NIGMS (1R35GM147706) to J.D.Y. We would like to thank Catherine Eichhorn and Daniel Herschlag for their thoughtful comments, which strengthened this paper.

## Contributions

J.D.Y. and C.J. designed the experiments. B.L., D.A., and K.N. performed the experiments. J.D.Y. performed the analysis with help from C.J. and D.H.S.D. J.D.Y. wrote the paper with the help of D.H.S.D. and all other authors.

## Methods

### Extracting isolated two-way junctions from high-resolution RNA structures

We extracted RNA structural motifs from high-resolution experimental structures using the RNA 3D Motif Atlas (http://rna.bgsu.edu/rna3dhub/nrlist) (44). This database provided non-redundant RNA structures determined by X-ray crystallography and cryo-EM with resolution better than 3.5 Å. Using DSSR (Dissecting the Spatial Structure of RNA) software (59), we identified all structural elements, including n-way junctions, two-way junctions, loops, and helices. At the same time, 3D Structures of Nucleic Acid-Protein complexes software (SNAP) (60) characterized any RNA-protein interactions present in these motifs. We focused on two-way junctions from these elements, applying filters to ensure structural stability when incorporated into designed constructs. We excluded junctions with more than two hydrogen bonds to non-junction nucleotides and removed those with extensive protein contacts or other tertiary interactions. This process yielded a set of two-way junctions that could maintain their native structural features when isolated. All motifs with known 3D structures used in this study are listed in **Supplemental Table S1**. The PDB of each structure used in this study is included in Fig Share (10.6084/m9.figshare.27880434).

### Selecting symmetric junction sequences that do not have 3D structures

We generated all 1x1 and 2x2 potential junction sequences with all possible flanking pairs (A-U, U-A, G-C, C-G, G-U, U-G). As we could not include all possible sequences, we applied selection creation. We focused on sequences that have the most As and Cs as possible as those that are sensitive to DMS modification.

### Design of RNA library of 7500 stable hairpin constructs

We designed RNA constructs containing 5-7 two-way junctions arranged in hairpin structures. Each construct included standardized primer sequences and a central hairpin loop, with junctions separated by 3 WC base pairs. Using ViennaRNA, we verified that each sequence folded into its intended structure with low ensemble defect scores (≤5). We filtered constructs to ensure lengths were within 10% of the minimum sequence length while not exceeding 150 nucleotides. We required a minimum hamming distance of 20 between all constructs to maintain sequence diversity. This design process continued iteratively until reaching 7,500 unique sequences, which were ordered as an Agilent oligo pool (sequences in **Supplemental Document: Sequences.xlsx**).

### PCR amplification of oligo pool to generate DNA templates

To generate double-stranded DNA templates for transcription, we amplified the oligo pool using PCR. The oligo pool was dissolved in 50 μL of 1x IDTE pH 8.0 buffer (IDT #11-05-01-13). The PCR reaction used forward (TTCTAATACGACTCACTATAGG) and reverse (GTTGTTGTTGTTGTTTCTTT) primers from IDT. Each 50 μL reaction contained 25 μL Q5 High-Fidelity DNA Polymerase (NEB #M0494S), 2 μL oligo pool, 2.5 μL each primer (diluted to 10 μM from 100 μM stock), and 18 μL RNase-free UltraPure water (ThermoFisher #10977015). PCR conditions were: 98°C for 30s, then 20 cycles of 98°C for 10s, 62°C for 15s, and 72°C for 15s, with final extension at 72°C for 5min. Products were separated on 2% agarose gel (150V, 1h) and purified using Zymoclean Gel DNA Recovery Kit (Genesee Scientific #11-301C).

### In vitro RNA synthesis and purification

RNA was transcribed in vitro using a 100 μL reaction containing: 10 μL 10x Transcription Buffer (400 mM Tris-HCl pH 8.0, 10 mM spermidine, 0.1% Triton X), 5 μL 50 mM DTT, 16 μL 25 mM NTPs, 8 μL 250 mM MgCl_2_, 4 μL T7 polymerase (NEB #M0251S), 24 μL template DNA (adjusted to 0.3 μM), and 33 μL RNase-free water. After 6 hour incubation at 37°C, DNA was removed with DNase I and RNA was purified using RNA Clean and Concentrator-5 kit (Genesee Scientific #R1014). Final RNA concentration was measured by nanodrop spectrophotometry and length was verified by 4% denaturing agarose gel electrophoresis (150V, 1h).

### DMS modification and library preparation for next-generation sequencing

DMS modification was performed on 10 pmol RNA in 5 μL RNase-free water. RNA was denatured (90°C, 4min), snap-cooled (4°C, 3min), then added to folding solution containing 16.5 μL buffer and 1 μL MgCl2 at optimized concentrations. To achieve final concentrations of 0.265 mM sodium cacodylate and 10 mM MgCl2, we combined 16.5 μL of 0.4 M sodium cacodylate with 1 μL of 250 mM MgCl_2_. RNA was folded at room temperature for 30 min. Meanwhile, DMS solution was prepared by mixing 15 μL DMS (Sigma-Aldrich #D186309) with 85 μL 100% ethanol (Decon Labs #2716). After folding, 2.5 μL DMS solution was added for 6 min, then quenched with 25 μL BME (ThermoFisher #125470010). Modified RNA was purified using RNA Clean & Concentrator-5 kit (Genesee Scientific #R1014), eluted in 7 μL RNase-free water, and quantified using Qubit RNA BR Assay Kit (ThermoFisher #Q10211) using 1 μL sample.

TGIRT-III reverse transcription was used to detect DMS modifications through mutation incorporation. The 12.1 μL reaction contained: 2.4 μL 5x TGIRT buffer (250 mM Tris-HCl pH 8.3, 375 mM KCl, 15 mM MgCl_2_), 1.2 μL 10 mM dNTPs, 0.6 μL 100 mM DTT, 0.5 μL TGIRT-III enzyme, 6.4 μL modified RNA (diluted to 0.25 μM), and 1 μL barcoded RTB primer (0.285 μM; **sequences in Supplemental Document: Sequences.xlsx**). After 2h incubation at 57°C, RNA was hydrolyzed by adding 5 μL 0.4 M NaOH, heating (90°C, 4min), and snap-cooling (4°C, 3min). The reaction was neutralized with 2.5 μL quench acid (1.43 M NaCl, 0.57 M HCl, 1.29 M sodium acetate; volume adjusted per batch). The cDNA was purified using Oligo Clean and Concentrator Kit (Genesee Scientific #11-380B), adding 30 μL RNase-free water before purification to reach 50 μL total volume. cDNA was eluted in 15 μL RNase-free water.

The cDNA library was amplified using PCR with forward primer AATGATACGGCGACCACCGAGATCTACACTCTTTCCCTACACGACGCTCTTCCG and reverse primer CAAGCAGAAGACGGCATACGAGATCGGTCTCGGCATTCCTGCTGAACCGCTCTTCCGATCTGGGCTTCGGCCC. Each 50 μL reaction contained 25 μL Q5 High-Fidelity DNA Polymerase (NEB #M0494S), 2.5 μL each primer, 2.0 μL purified cDNA, and 18 μL RNase-free water. PCR conditions were: 98°C for 30s, then 16 cycles of 98°C for 10s, 62°C for 15s, and 72°C for 15s, with final extension at 72°C for 5min. Products were separated on 2% E-gel EX Agarose Gel (ThermoFisher #G401002) using E-Gel Power Snap Plus system (ThermoFisher #G9301) for 10min. Correct size bands were excised and purified using Zymoclean Gel DNA Recovery Kit (Genesee Scientific #11-301C). Final library concentration was measured using Qubit 1X dsDNA High Sensitivity Assay Kit (ThermoFisher #Q33230).

### Generation of DMS reactivity from DMS-MaPseq sequencing data

The sequencing was conducted using Novaseq 6000. The sequencing run was initially demultiplexed using the RTB barcodes inserted during the RT process, utilizing the novobarcode software (https://www.novocraft.com/documentation/novobarcode/demultiplexing-barcodedindexed-reads-with-novobarcode/).

`novobarcode -b rtb_barcodes.fa -f test_R1_001.fastq test_R2_001.fastq`

An example for rtb_barcodes.fa would be;

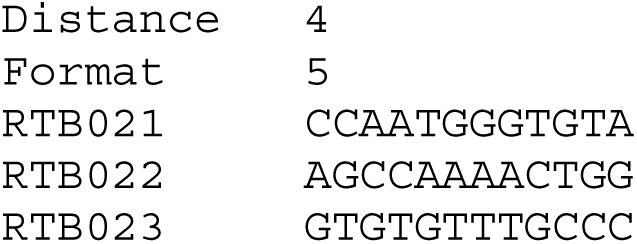

The `Distance` refers to the variation in base pairs between a barcode and a permissible read when three barcodes are provided. The format specifies that the barcode will be located at the 5′ end of read 1. These demultiplexed fastq files are available on Sequencing Read Archive accession PRJNA1188187.

Processing the demultiplexed fastq files into the mutation fractions was performed by the rna-map software (https://github.com/YesselmanLab/rna_map) (61). With the following command for each replicate

`rna-map -fa <FASTA file> -fq1 <R2 fastq file> -fq2 <R1 fastq file> -- dot-bracket <CSV file>`

Where `<FASTA file>` is the path to the fasta file with all library DNA sequences without T7 promoters. `<R2 fastq file>` is the path to the R2 fastq file obtained from demultiplexing.

`<R1 fastq file>` is the path to the R1 fastq file obtained from demultiplexing. `<CSV file>` is an optional file that contains the name, RNA sequence and structure in dot bracket notation for each sequence in the library. `rna-map` will generate a ‘mutation_histos.p’ file that will be used for the next analysis steps.

### Processing DMS reactivity for motif and residue analysis

To ensure high quality data, we filtered our initial dataset of 7,500 sequences, removing any sequences with fewer than 2,000 reads or signal-to-noise ratios below 4. This filtering step eliminated 17 sequences. We also excluded individual reactivity measurements with z-scores exceeding 3 (approximately 2,424 datapoints, ∼1% of total data). These outliers were predominantly found in flanking Watson-Crick pairs from a small subset of motif sequences (**Supplemental Figure S16, Supplemental Table S4**), suggesting sequence-dependent alternative conformations in these cases.

Each construct sequence was parsed into motifs based on secondary structure and sequence. We defined the first flanking pair as the Watson-Crick pair directly neighboring non-canonical interactions, which remained constant for motifs derived from 3D structures. The second flanking pair was defined as the next Watson-Crick pair beyond the first flanking pair, which varied between constructs. To ensure consistent analysis, we standardized motif sequences by always placing the longer strand first (e.g., "GG&CAG" became "CAG&GG").

For each unique motif sequence, we calculated average reactivity values, standard deviations, and coefficients of variation. We then cross-referenced motifs with their corresponding PDB structures to extract detailed structural information. For each nucleotide in each motif, we collected nucleotide residue types (e.g., Flanking-WC, non-WC), sequential position, reactivity data, and pairing information. The residue data were expanded to include neighboring residue types (purine or pyrimidine) and stacking information between nucleotides. After log-transforming residue reactivity data, we mapped each residue to its corresponding PDB file and retrieved additional structural parameters (e.g., B-factors) for residues in the PDB files.

### Logistic Regression for Predicting Watson-Crick Base Pairs from Reactivity Data

This method employs logistic regression to predict the probability that a base pair is either WC or NON-WC based on the natural logarithmic reactivity data of nucleotides. Logistic regression is a binary classification technique that applies a sigmoid function to the input data, producing a probability score between 0 and 1, where values closer to 1 indicate a higher likelihood of the base pair being WC. In this method, the base pair type is first transformed into a binary variable, with WC encoded as 1 and non-WC as 0. The logistic regression model is then trained on the natural logarithmic reactivity data, learning the relationship between reactivity and base pair type. Once trained, the model computes the probability that each base pair is WC.

### Calculating the structural features for nucleotides in motifs that have PDB files

We analyzed the structural parameters of RNA junctions using the 3DNA software package

(60). For each PDB file, the `find_pair` and `analyze` commands were executed to generate base-pair parameters.

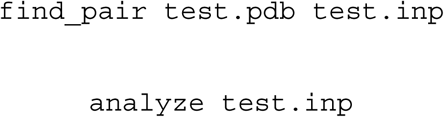

From the 3DNA output, we extracted detailed base-pair parameters including classification (WC or non-WC), residue numbers, and geometric measurements (shear, stretch, stagger, buckle, propeller, and opening angles). To assess structural deviations, we calculated root-mean-square deviations (RMSDs) by aligning base pairs to ideal PDB conformations using the Kabsch algorithm. We then characterized non-canonical base pairs using the Leontis-Westhof classification system (62), manually comparing each structure to exemplars from the RNA Basepair Catalog (63). This manual approach avoided misclassification errors we observed with automated methods. Using the Biopython PandasPdb module, we extracted atomic coordinates to calculate pairwise distances between all atoms in residues of interest. Finally, we calculated solvent accessible surface area (SASA) for specific atoms using the freesasa package.

## References

1 Ban N, Nissen P, Hansen J, Moore PB, Steitz TA. The complete atomic structure of the large ribosomal subunit at 2.4 Å resolution. Science. 2000;289(5481):905–20.

2 Jiang J, Wang Y, Susac L, Chan H, Basu R, Zhou ZH, et al. Structure of Telomerase with Telomeric DNA. Cell. 2018;173(5):1179–90 e13.

3 Schmitzová J, Cretu C, Dienemann C, Urlaub H, Pena V. Structural basis of catalytic activation in human splicing. Nature. 2023;617(7962):842-+.

4 Herschlag D, Bonilla S, Bisaria N. The Story of RNA Folding, as Told in Epochs. Cold Spring Harb Perspect Biol. 2018;10(10).

5 Kavita K, Breaker RR. Discovering riboswitches: the past and the future. Trends in Biochemical Sciences. 2023;48(2):119–41.

6 Micura R, Höbartner C. Fundamental studies of functional nucleic acids: aptamers, riboswitches, ribozymes and DNAzymes. Chemical Society Reviews. 2020;49(20):7331–53.

7 Steitz TA. A structural understanding of the dynamic ribosome machine. Nature Reviews Molecular Cell Biology. 2008;9(3):242–53.

8 Tholen J, Galej WP. Structural studies of the spliceosome: Bridging the gaps. Current Opinion in Structural Biology. 2022;77.

9 Ganser LR, Kelly ML, Herschlag D, Al-Hashimi HM. The roles of structural dynamics in the cellular functions of RNAs. Nature Reviews Molecular Cell Biology. 2019;20(8):474–89.

10 Vicens Q, Kieft JS. Thoughts on how to think (and talk) about RNA structure. Proc Natl Acad Sci U S A. 2022;119(17):e2112677119.

11 Cordero P, Das R. Rich RNA Structure Landscapes Revealed by Mutate-and-Map Analysis. Plos Comput Biol. 2015;11(11).

12 Ehrhardt JE, Weeks KM. Time-Resolved, Single-Molecule, Correlated Chemical Probing of RNA. J Am Chem Soc. 2020;142(44):18735–40.

13 Lange B, Gil RG, Anderson GS, Yesselman JD. High-throughput determination of RNA tertiary contact thermodynamics by quantitative DMS chemical mapping. Nucleic Acids Res. 2024;52(16):9953–65.

14 Martin S, Blankenship C, Rausch JW, Sztuba-Solinska J. Using SHAPE-MaP to probe small molecule-RNA interactions. Methods. 2019;167:105–16.

15 Mustoe AM, Lama NN, Irving PS, Olson SW, Weeks KM. RNA base-pairing complexity in living cells visualized by correlated chemical probing. Proc Natl Acad Sci U S A. 2019;116(49):24574–82.

16 Mustoe AM, Weidmann CA, Weeks KM. "Single-Molecule Correlated Chemical Probing: A Revolution in RNA Structure Analysis" (vol 56, pg 763, 2023). Accounts of Chemical Research. 2023;56(12):1684-.

17 Olson SW, Turner AW, Arney JW, Saleem I, Weidmann CA, Margolis DM, et al. Discovery of a large-scale, cell-state-responsive allosteric switch in the 7SK RNA using DANCE-MaP. Mol Cell. 2022;82(9):1708–23 e10.

18 Strobel EJ, Cheng LY, Berman KE, Carlson PD, Lucks JB. A ligand-gated strand displacement mechanism for ZTP riboswitch transcription control. Nat Chem Biol. 2019;15(11):1067-+.

19 Strobel EJ, Yu AM, Lucks JB. High-throughput determination of RNA structures. Nat Rev Genet. 2018;19(10):615–34.

20 Tijerina P, Mohr S, Russell R. DMS footprinting of structured RNAs and RNA-protein complexes. Nat Protoc. 2007;2(10):2608–23.

21 Tomezsko P, Swaminathan H, Rouskin S. Viral RNA structure analysis using DMS-MaPseq. Methods. 2020;183:68–75.

22 Tomezsko P, Swaminathan H, Rouskin S. DMS-MaPseq for Genome-Wide or Targeted RNA Structure Probing In Vitro and In Vivo. Methods Mol Biol. 2021;2254:219–38.

23 Weidmann CA, Mustoe AM, Jariwala PB, Calabrese JM, Weeks KM. Analysis of RNA-protein networks with RNP-MaP defines functional hubs on RNA. Nat Biotechnol. 2021;39(3):347–56.

24 Zubradt M, Gupta P, Persad S, Lambowitz AM, Weissman JS, Rouskin S. DMS-MaPseq for genome-wide or targeted RNA structure probing in vivo. Nat Methods. 2017;14(1):75–82.

25 Homan PJ, Favorov OV, Lavender CA, Kursun O, Ge XY, Busan S, et al. Single-molecule correlated chemical probing of RNA. P Natl Acad Sci USA. 2014;111(38):13858–63.

26 Cordero P, Kladwang W, VanLang CC, Das R. Quantitative Dimethyl Sulfate Mapping for Automated RNA Secondary Structure Inference. Biochemistry-Us. 2012;51(36):7037–9.

27 Ding YL, Tang Y, Kwok CK, Zhang Y, Bevilacqua PC, Assmann SM. genome-wide profiling of RNA secondary structure reveals novel regulatory features. Nature. 2014;505(7485):696-+.

28 Hajdin CE, Bellaousov S, Huggins W, Leonard CW, Mathews DH, Weeks KM. Accurate SHAPE-directed RNA secondary structure modeling, including pseudoknots. P Natl Acad Sci USA. 2013;110(14):5498–503.

29 Mathews DH, Disney MD, Childs JL, Schroeder SJ, Zuker M, Turner DH. Incorporating chemical modification constraints into a dynamic programming algorithm for prediction of RNA secondary structure. P Natl Acad Sci USA. 2004;101(19):7287–92.

30 Wu Y, Shi BB, Ding XQ, Liu T, Hu XH, Yip KY, et al. Improved prediction of RNA secondary structure by integrating the free energy model with restraints derived from experimental probing data. Nucleic Acids Res. 2015;43(15):7247–59.

31 Allan MF, Aruda J, Plung JS, Grote SL, des Taillades YJM, de Lajarte AA, et al. Discovery and Quantification of Long-Range RNA Base Pairs in Coronavirus Genomes with SEARCH-MaP and SEISMIC-RNA. Res Sq. 2024.

32 Lan TCT, Allan MF, Malsick LE, Woo JZ, Zhu C, Zhang FR, et al. Secondary structural ensembles of the SARS-CoV-2 RNA genome in infected cells. Nat Commun. 2022;13(1).

33 Morandi E, Manfredonia I, Simon LM, Anselmi F, van Hemert MJ, Oliviero S, et al. Genome-scale deconvolution of RNA structure ensembles. Nature Methods. 2021;18(3):249-+.

34 Olson SW, Turner AMW, Arney JW, Saleem I, Weidmann CA, Margolis DM, et al. Discovery of a large-scale, cell-state-responsive allosteric switch in the 7SK RNA using DANCE-MaP. Mol Cell. 2022;82(9):1708-+.

35 Tomezsko PJ, Corbin VDA, Gupta P, Swaminathan H, Glasgow M, Persad S, et al. Determination of RNA structural diversity and its role in HIV-1 RNA splicing (vol 582, pg 438, 2020). Nature. 2020;588(7837):E16–E.

36 Cordero P, Kladwang W, VanLang CC, Das R. Quantitative dimethyl sulfate mapping for automated RNA secondary structure inference. Biochemistry. 2012;51(36):7037–9.

37 Das R, Karanicolas J, Baker D. Atomic accuracy in predicting and designing noncanonical RNA structure. Nat Methods. 2010;7(4):291–4.

38 Muth GW, Ortoleva-Donnelly L, Strobel SA. A single adenosine with a neutral pKa in the ribosomal peptidyl transferase center. Science. 2000;289(5481):947-50.

39 Wells SE, Hughes JM, Igel AH, Ares M, Jr. Use of dimethyl sulfate to probe RNA structure in vivo. Methods Enzymol. 2000;318:479–93.

40 Noller HF, Woese CR. Secondary structure of 16S ribosomal RNA. Science. 1981;212(4493):403-11.

41 Ehresmann C, Baudin F, Mougel M, Romby P, Ebel JP, Ehresmann B. Probing the structure of RNAs in solution. Nucleic Acids Res. 1987;15(22):9109–28.

42 Romby P, Westhof E, Toukifimpa R, Mache R, Ebel JP, Ehresmann C, et al. Higher order structure of chloroplastic 5S ribosomal RNA from spinach. Biochemistry. 1988;27(13):4721–30.

43 Das R, Baker D. Automated de novo prediction of native-like RNA tertiary structures. Proc Natl Acad Sci U S A. 2007;104(37):14664–9.

44 Leontis NB, Zirbel CL. Nonredundant 3D Structure Datasets for RNA Knowledge Extraction and Benchmarking. In: Leontis N, Westhof E, editors. RNA 3D Structure Analysis and Prediction. Berlin, Heidelberg: Springer Berlin Heidelberg; 2012. p. 281-98.

45 Leontis NB, Lescoute A, Westhof E. The building blocks and motifs of RNA architecture. Curr Opin Struct Biol. 2006;16(3):279–87.

46 Jasinski D, Haque F, Binzel DW, Guo P. Advancement of the Emerging Field of RNA Nanotechnology. ACS Nano. 2017;11(2):1142–64.

47 Yesselman JD, Eiler D, Carlson ED, Gotrik MR, d’Aquino AE, Ooms AN, et al. Computational design of three-dimensional RNA structure and function. Nat Nanotechnol. 2019;14(9):866–73.

48 Ellington AD, Szostak JW. In vitro selection of RNA molecules that bind specific ligands. Nature. 1990;346(6287):818-22.

49 Ferre-D’amare A R, Rupert PB. The hairpin ribozyme: from crystal structure to function. Biochem Soc Trans. 2002;30(Pt 6):1105–9.

50 Geng A, Roy R, Al-Hashimi HM. Conformational penalties: New insights into nucleic acid recognition. Curr Opin Struct Biol. 2024;89:102949.

51 Huang L, Lilley DMJ. The kink-turn in the structural biology of RNA. Q Rev Biophys. 2018;51:e5.

52 Szewczak AA, Moore PB. The sarcin/ricin loop, a modular RNA. J Mol Biol. 1995;247(1):81–98.

53 Bailor MH, Sun XY, Al-Hashimi HM. Topology Links RNA Secondary Structure with Global Conformation, Dynamics, and Adaptation. Science. 2010;327(5962):202-6.

54 Al-Hashimi HM, Gosser Y, Gorin A, Hu W, Majumdar A, Patel DJ. Concerted motions in HIV-1 TAR RNA may allow access to bound state conformations: RNA dynamics from NMR residual dipolar couplings. J Mol Biol. 2002;315(2):95–102.

55 Faison EM, Nallathambi A, Zhang Q. Characterizing Protonation-Coupled Conformational Ensembles in RNA via pH-Differential Mutational Profiling with DMS Probing. J Am Chem Soc. 2023;145(34):18773–7.

56 Kotar A, Ma S, Keane SC. pH dependence of C*A, G*A and A*A mismatches in the stem of precursor microRNA-31. Biophys Chem. 2022;283:106763.

57 Legault P, Pardi A. In-Situ Probing of Adenine Protonation in Rna by C-13 Nmr. Journal of the American Chemical Society. 1994;116(18):8390–1.

58 Wilcox JL, Bevilacqua PC. pKa shifting in double-stranded RNA is highly dependent upon nearest neighbors and bulge positioning. Biochemistry. 2013;52(42):7470–6.

59 Lu XJ, Bussemaker HJ, Olson WK. DSSR: an integrated software tool for dissecting the spatial structure of RNA. Nucleic Acids Res. 2015;43(21).

60 Lu XJ, Olson WK. 3DNA: a versatile, integrated software system for the analysis, rebuilding and visualization of three-dimensional nucleic-acid structures. Nature Protocols. 2008;3(7):1213–27.

61 Jurich CP, Brivanlou A, Rouskin S, Yesselman JD. Web-based platform for analysis of RNA folding from high throughput chemical probing data. Nucleic Acids Res. 2022;50(W1):W266–W71.

62 Leontis NB, Westhof E. Geometric nomenclature and classification of RNA base pairs. Rna. 2001;7(4):499–512.

63 Lawson CL, Berman HM, Chen L, Vallat B, Zirbel CL. The Nucleic Acid Knowledgebase: a new portal for 3D structural information about nucleic acids. Nucleic Acids Res. 2023.

