## Supplemental Material for "Characterizing 3D RNA structural features from DMS reactivity"

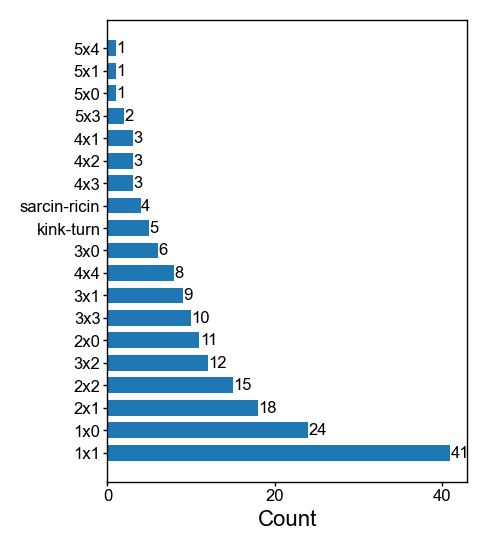

### Supplemental Figure S1: Distribution of two-way junctions

The horizontal bar plot shows the counts of different two-way junctions in the library. The motifs are sorted by their frequency.

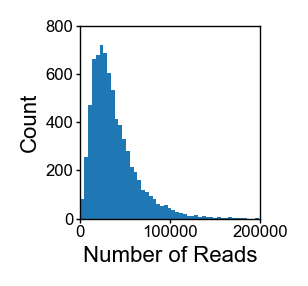

### Supplemental Figure S2: Number of reads per construct

This histogram represents the distribution of the number of reads per construct. Most of the distribution falls between 0 and 100,000 reads, with the highest frequency observed at lower read counts. The distribution exhibits a long tail, indicating a small subset of constructs with higher read counts.

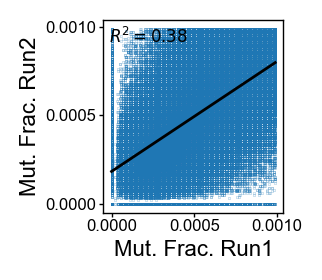

### Supplemental Figure S3: Reproducibility of DMS measurements below 0.001 across independent experiments.

The correlation plot illustrates the R² value between two independent experiments for DMS reactivity values below 0.001. When considering the entire data range, the correlation was strong (R² = 0.99**, Figure 1C**), but it significantly decreased to 0.38 for DMS reactivity values below 0.001**.**

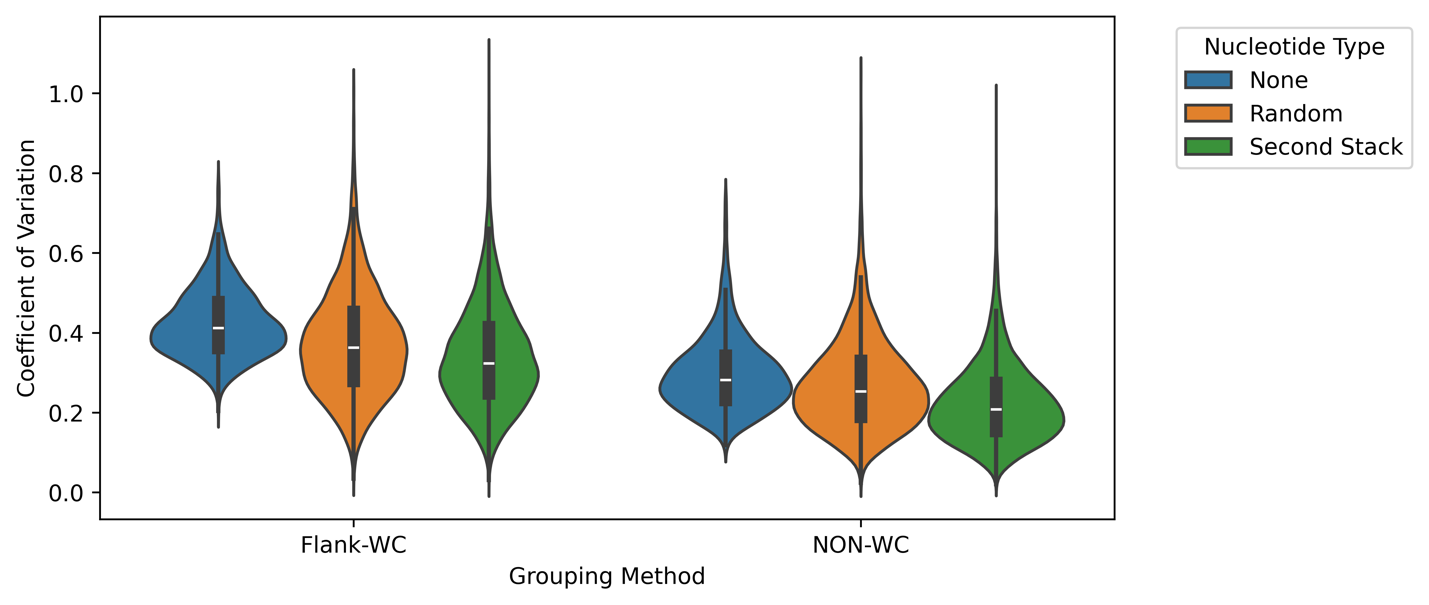

### Supplemental Figure S4: Comparison of variability between second flanking pair grouping and random grouping

Violin plots illustrating the coefficient of variation (CV) across different grouping methods: no grouping (None), random grouping into similarly sized groups (Random), and grouping based on second base pair stacking (Second Stack). The results show minimal differences in variability between random grouping and second base pair stacking, suggesting that the grouping method has little impact on the overall variability in CV.

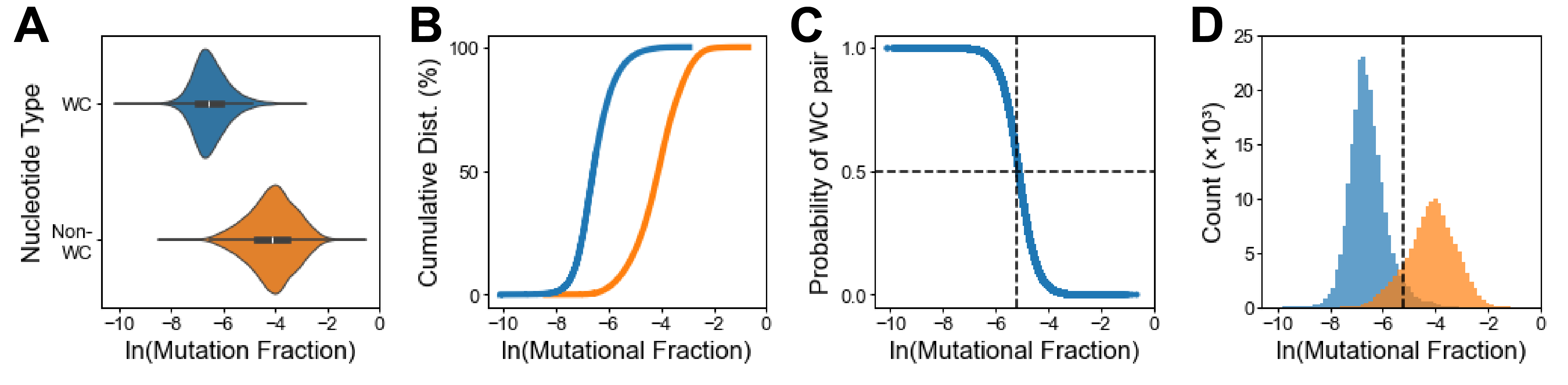

### Supplemental Figure S5: Quantitative analysis of DMS reactivity in WC and non-WC nucleotides

Remaking Figure 2 including all WC pairs not just flanking pairs, the results are similar to just including flanking WC pairs (A) Reactivity distribution of Non-WC and WC nucleotides. (B) Cumulative reactivity distributions comparing WC paired versus non-WC nucleotides as a function of the natural log of the mutational histogram. (C) Logistic regression analysis establishing the probability of WC pairing based on DMS reactivity. The horizontal dashed line marks the 50% probability threshold, corresponding to a natural log mutation fraction of -5.19 (mutation fraction = 0.0056). (D) Distribution of nucleotides relative to the 50% probability threshold.

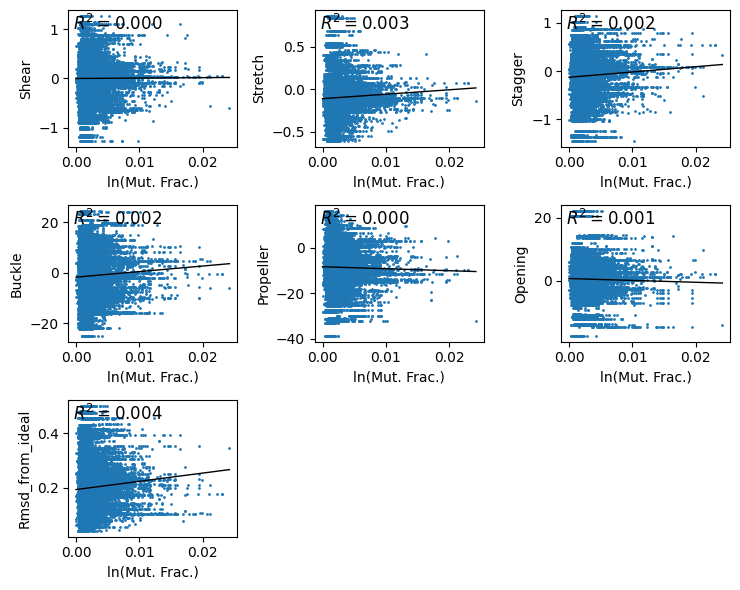

### Supplemental Figure S6: Quantitative analysis of reactivity of flanking WC pairs to base pair parameters

Scatter plots illustrating the correlation between natural logarithm of DMS reactivity and the six base pair parameters: shear, stretch, stagger, buckle, propeller, and opening. R2 values indicate no correlation between the reactivity and the parameters.

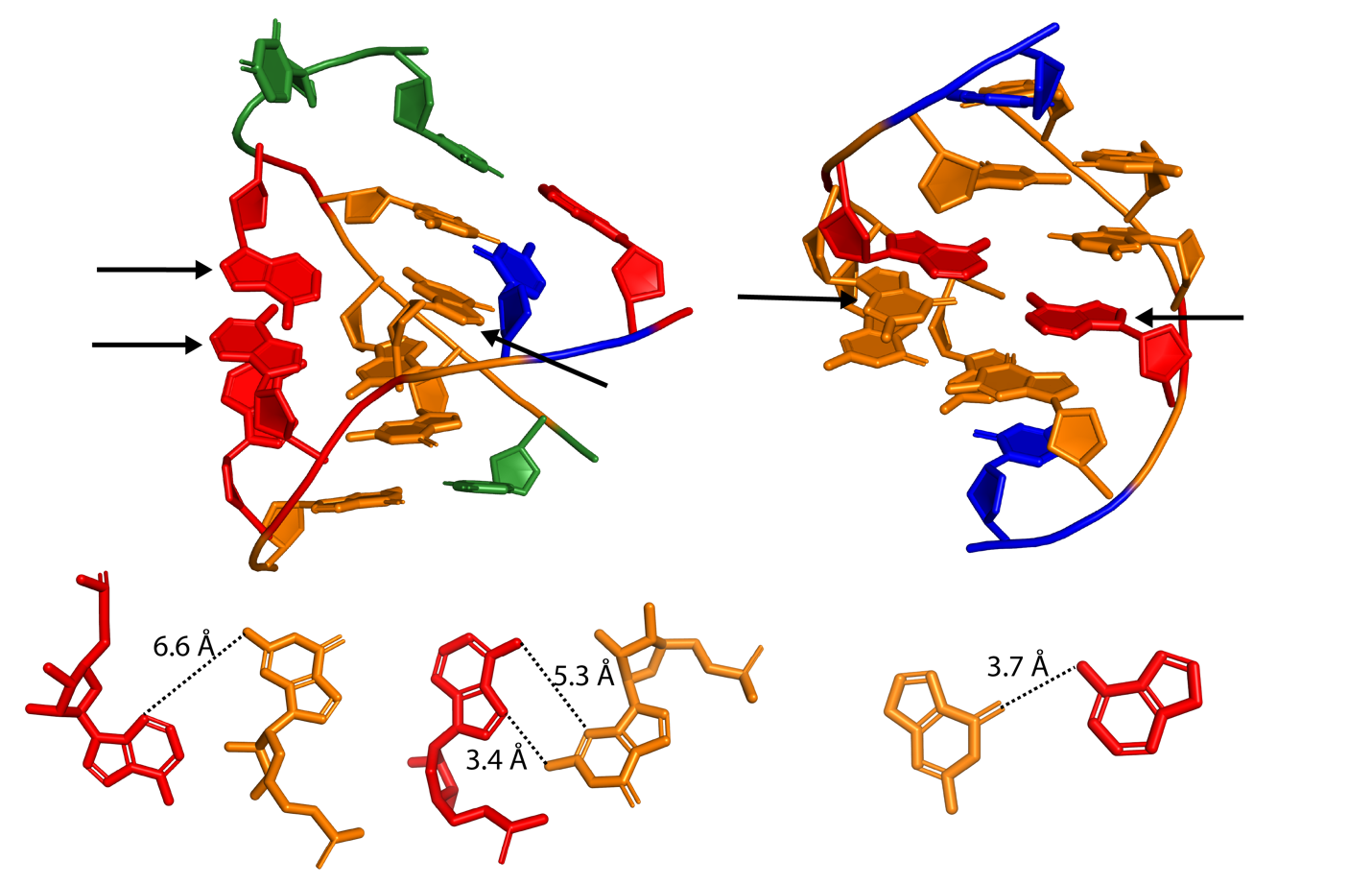

### Supplemental Figure S7: Purines’ preference for stacking interactions over hydrogen bonding in purine-rich environments

This figure depicts the structural preferences of purines in purine-rich environments, where stacking interactions are favored over hydrogen bonding. The top panels show molecular arrangements highlighting purine stacking interactions with arrows showing the regions of interest. The bottom panels present the measured distances between purine bases, demonstrating that these distances exceed the typical hydrogen bond length (< 3.3 Å), emphasizing the dominance of stacking interactions in these configurations. These are two examples from the library, representing instances of purine stacking behavior.

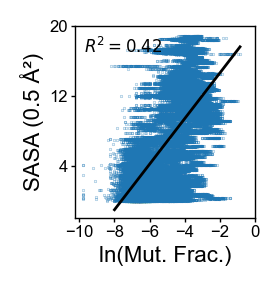

### Supplemental Figure S8: Correlation between solvent accessible surface area (SASA) and mutation fraction

This scatter plot depicts the relationship between the solvent accessible surface area (measured at 0.5 Å probe radius) and the natural logarithm of the mutation fraction. The linear regression line demonstrates a moderate positive correlation, with an R² value of 0.42.

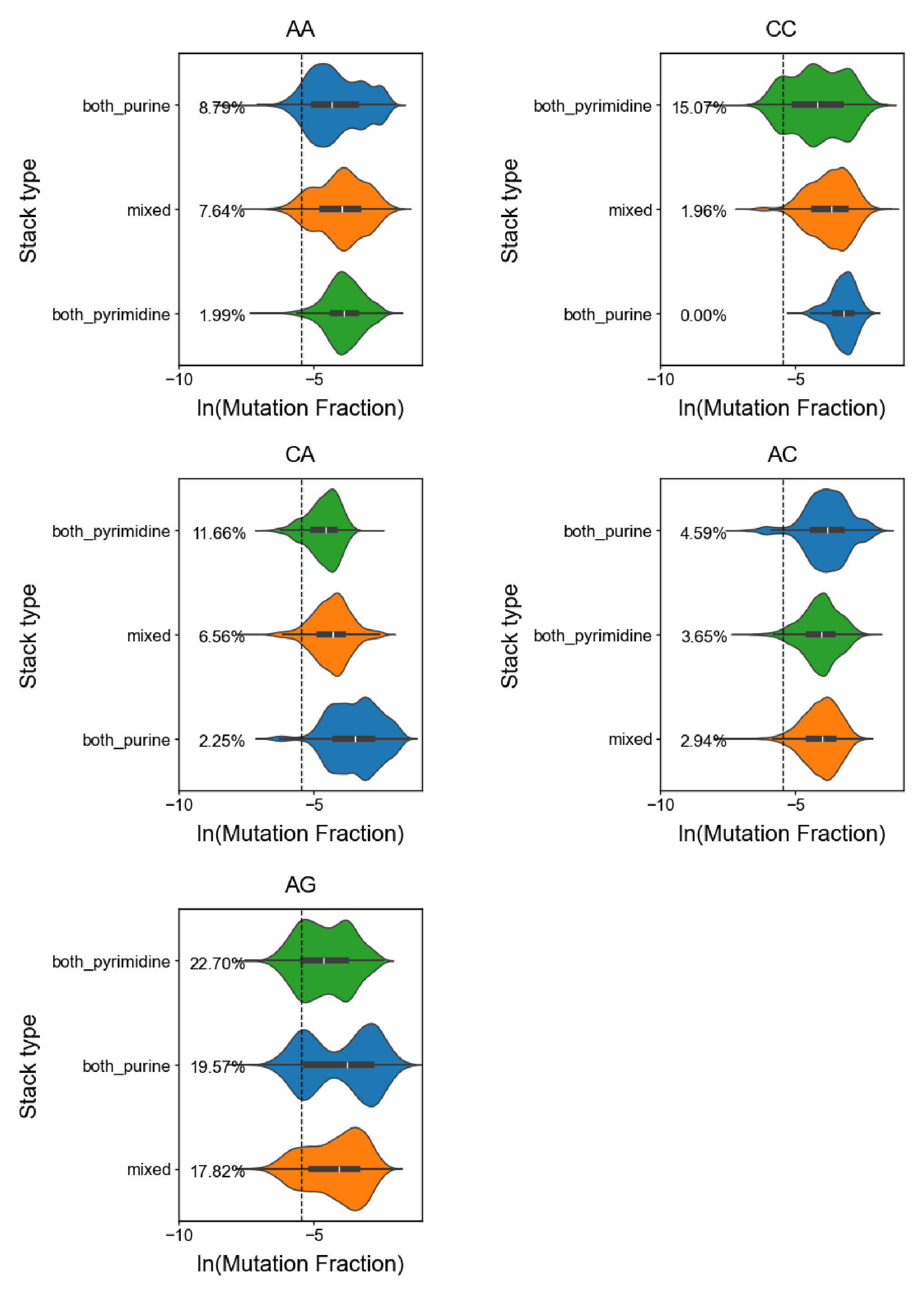

### Supplemental Figure S9: Impact of neighboring sequences on mismatches

Violin plots illustrating the impact of neighboring nucleotide sequences on mismatch mutation fractions. Each plot represents the distribution of ln(Mutation Fraction) for specific neighboring sequence contexts (A-A, C-C, C-A, A-C, and A-G), categorized into stack types: both_purine, mixed, and both_pyrimidine. The horizontal dashed line indicates the 50% probability threshold (-5.45). The percentages indicate the proportion of data falling ln(Mutation Fraction) < -5.45.

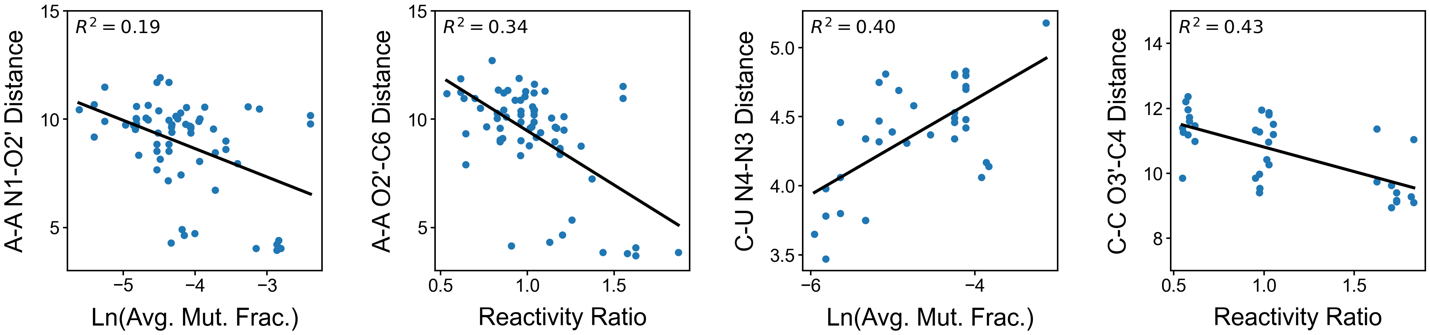

### Supplemental Figure S10: Weaker correlations between non-canonical pairs and DMS reactivity

Scatter plots showing weaker correlations between DMS reactivity and specific atomic distances in non-canonical base pairs. Each plot represents a distinct atomic pair (e.g., A N1–O2’, A O2–C6, C-U N4–N3, and C-C O3’–C4), with the associated R^2^ values indicating the strength of the linear correlation. Reactivity is expressed as either the natural logarithm of average mutational fraction or the reactivity ratio of the two residues in the mismatch.

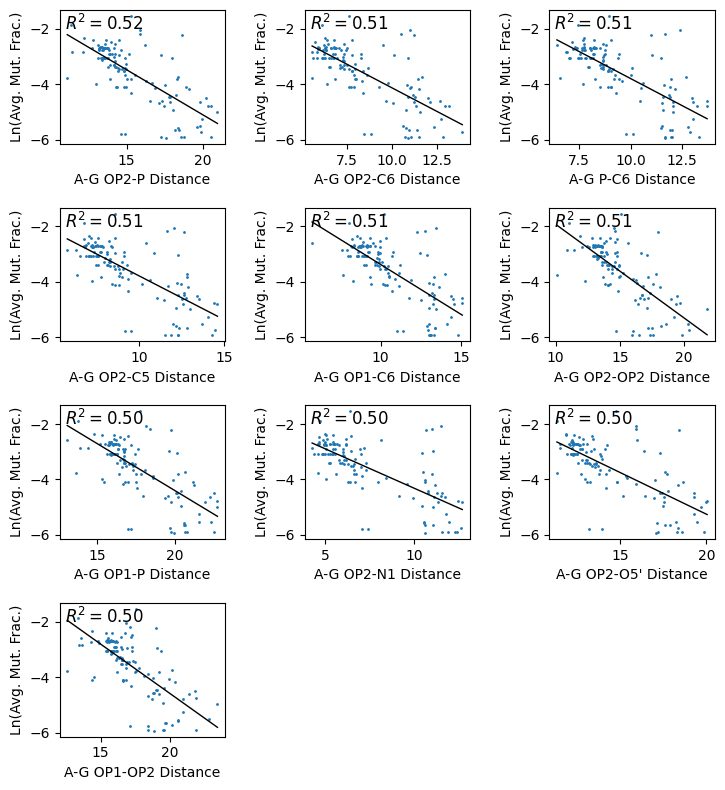

### Supplemental Figure S11: Correlation plots for distance and reactivity for A-G pairs

Scatter plots showing the top 10 correlations between atomic distances in A-G pairs and the natural logarithm of DMS reactivity of the A. Each plot represents a specific atomic pair distance (e.g., OP2-P, OP2-C6, OP1-C6), with the corresponding R² values indicating the strength of the correlation.

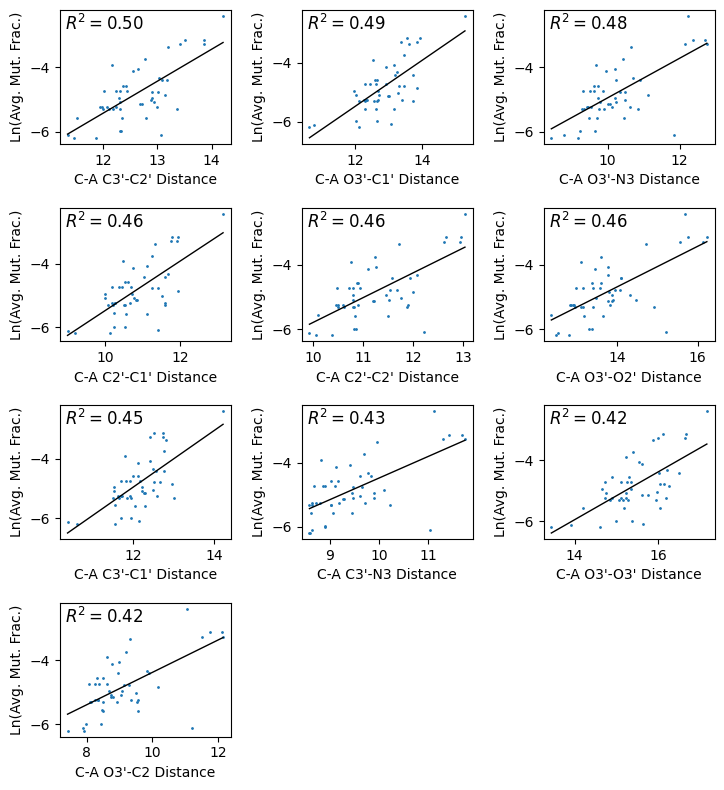

### Supplemental Figure S12: Correlation plots for distance and reactivity for C-A pairs

Scatter plots showing the top 10 correlations between atomic distances of C in C-A pairs and the natural logarithm of DMS reactivity. Each plot represents a specific atomic pair distance (e.g., C3′-C2′, O3′-C1′, O3′-N3), with the corresponding R² values indicating the strength of the correlation.

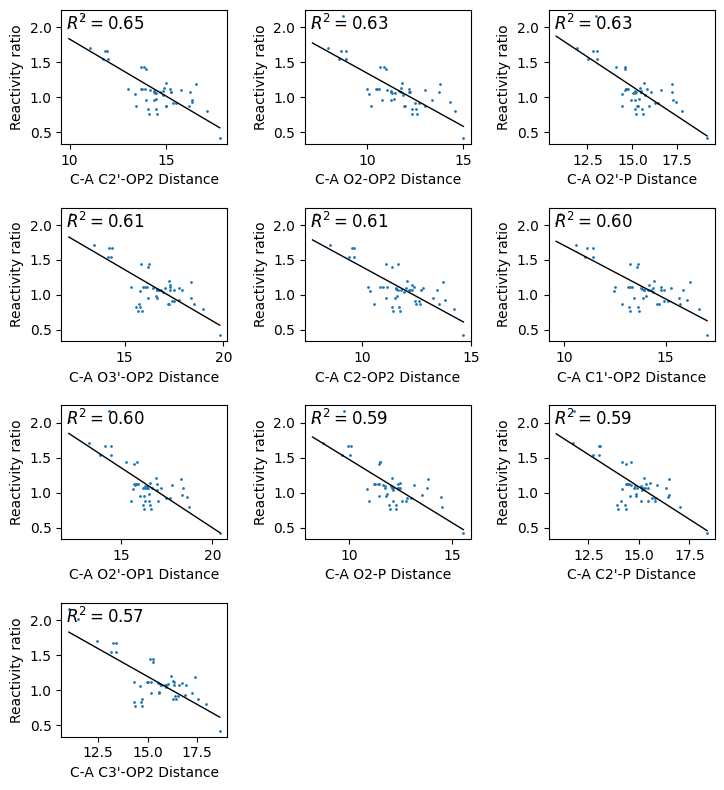

### Supplemental Figure S13: Correlation plots for distance and reactivity ratio for C-A pairs

Scatter plots showing the top 10 correlations between atomic distances of C in C-A pairs and the ratio of the natural logarithm of DMS reactivates. Each plot represents a specific atomic pair distance (e.g., C2′-OP2, O2-OP2, O2′-P), with the corresponding R² values indicating the strength of the correlation.

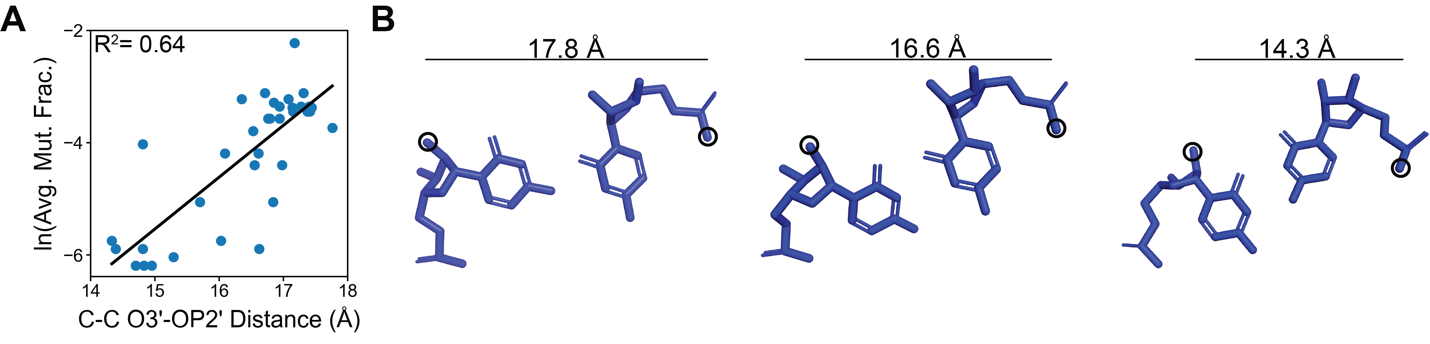

### Supplemental Figure S14: Correlation between cytosine reactivity and atomic distance for C-C mismatches.

(A) Scatter plot illustrating the correlation between cytosine reactivity, expressed as ln(Avg. Mut. Frac.), and the C-C O3’–OP2’ distance. The R^2^ value of 0.64 shows a strong correlation. (B) Representative structural models of C-C mismatches with varying O3’–OP2’ distances (17.8 Å, 16.6 Å, and 14.3 Å), showing the spatial arrangement of cytosine residues.

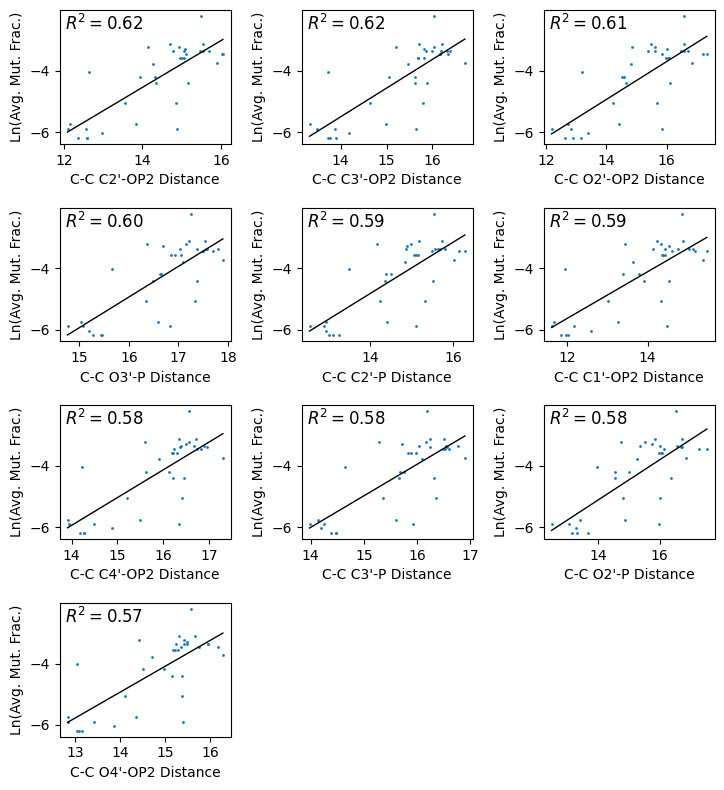

### Supplemental Figure S15: Correlation plots for distance and reactivity for C-C pairs

Scatter plots showing the top 10 correlations between interatomic distances in C-C pairs and the natural logarithm of DMS reactivity. Each plot represents a specific atomic pair distance (e.g., C2′-OP2, C3′-OP2, O2′-OP2), with the corresponding R² values indicating the strength of the correlation.

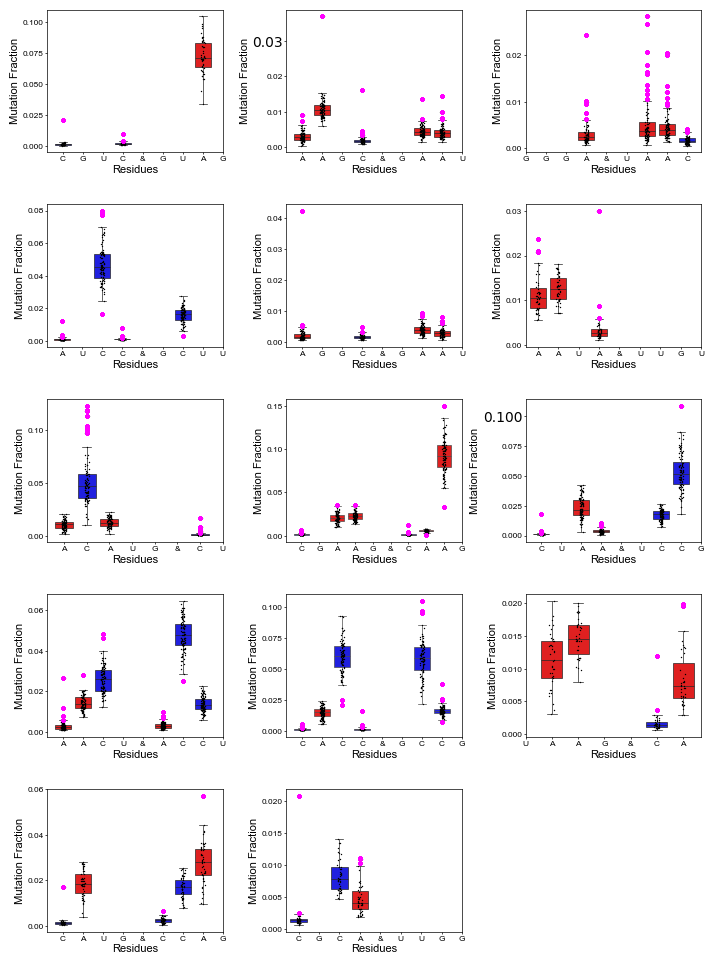

### Supplementary Figure S16: DMS reactivity patterns for motifs highlighting outliers

Box plots depict the reactivity ranges for residues in motifs that has outliers. The outlier reactivity values are highlighted in magenta.

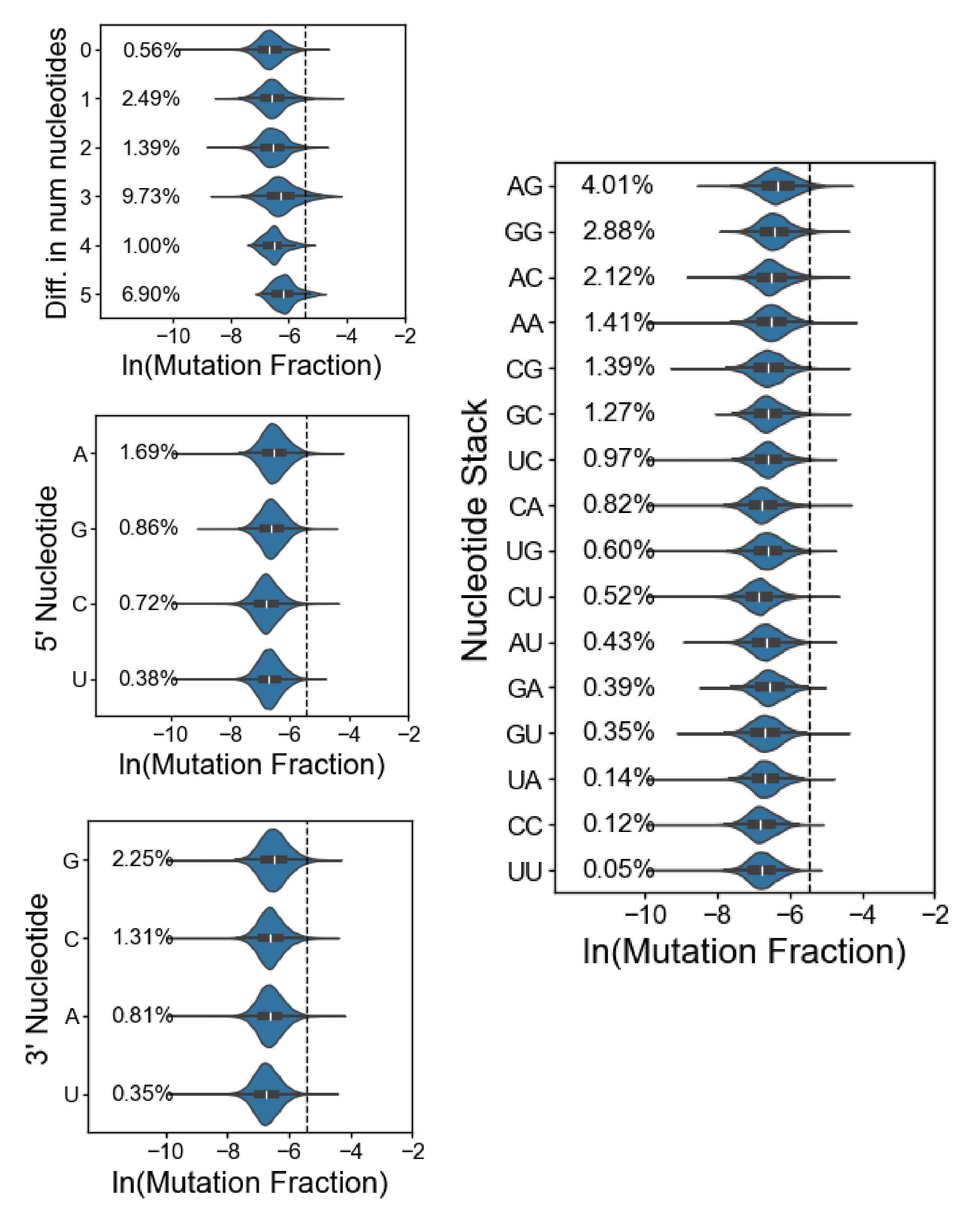

### Supplementary Figure S17: Reactive flanking base pairs for C

Remaking figures 3D-3G for cytosines. Similar trends were observed for C here as mentioned in **Figure 3D-G**.

| Category | Percent below 2 Å | Count |
| --- | --- | --- |
| A in A-A | 23.29 | 992 |
| A in A-C | 62.43 | 668 |
| A in A-G | 69.25 | 374 |
| C in C-A | 51.66 | 664 |
| C in C-C | 12.06 | 506 |
| C in C-U | 0.00 | 521 |

### Supplemental Table S3: Residues under 2 $\boldsymbol{Å}$ for solvent accessible surface area (SASA).

Table shows the number of residues and their corresponding percentages that are under 2 Å for solvent accessible surface area (SASA) for each base pair mismatch category.

.

| Motif | Nuc | Pos | Avg | SD | CV | Loop | Type | ln(DMS) |
| --- | --- | --- | --- | --- | --- | --- | --- | --- |
| CGUC_GUAG | C | 3 | 0.002 | 0.003 | 1.451 | 2,2 | WC Paired | -6.258 |
| AAGC_GAAU | C | 6 | 0.002 | 0.002 | 1.175 | 2,2 | WC Paired | -6.297 |
| UAAC_GGGA | A | 4 | 0.005 | 0.005 | 1.087 | 2,2 | Non WC Paired | -5.382 |
| ACGG_CAAU | C | 11 | 0.002 | 0.002 | 1.266 | 2,2 | WC Paired | -6.328 |
| AUCC_GCUU | A | 3 | 0.001 | 0.002 | 1.516 | 2,2 | WC Paired | -6.830 |
| GAAU_AGGC | A | 11 | 0.004 | 0.006 | 1.616 | 2,2 | WC Paired | -5.634 |
| UUGU_AAUA | A | 14 | 0.004 | 0.004 | 1.153 | 2,2 | WC Paired | -5.628 |
| CU_ACAUG | C | 3 | 0.002 | 0.003 | 1.171 | 0,3 | WC Paired | -6.052 |
| CGAAG_CAAG | C | 12 | 0.001 | 0.002 | 1.113 | 3,2 | WC Paired | -6.510 |
| UCCG_CUAA | C | 11 | 0.001 | 0.002 | 1.613 | 2,2 | WC Paired | -6.524 |
| AACU_ACCU | A | 3 | 0.003 | 0.004 | 1.371 | 2,2 | WC Paired | -5.943 |
| CACC_GCCG | C | 6 | 0.002 | 0.002 | 1.258 | 2,2 | WC Paired | -6.366 |
| UAAG_CA | C | 11 | 0.002 | 0.002 | 1.169 | 2,0 | WC Paired | -6.232 |
| CCAG_CAUG | C | 11 | 0.002 | 0.002 | 1.353 | 2,2 | WC Paired | -6.389 |
| UUGG_CGCA | C | 11 | 0.002 | 0.003 | 1.520 | 2,2 | WC Paired | -6.288 |

### Supplemental Table S4: Outliers of the coefficient of variation (CV).

This table presents outlier DMS reactivity values for nucleotides across different motifs, providing additional details about each outlier, including the coefficient of variation (CV), nucleotide type, position, loop, type, *etc.*
